# The RSK-RPS6 Axis Controls the Preconditioning Effect and Induces Spinal Cord Regeneration

**DOI:** 10.1101/2022.08.16.502928

**Authors:** Charlotte Decourt, Julia Schaeffer, Beatrice Blot, Antoine Paccard, Blandine Excoffier, Mario Pende, Homaira Nawabi, Stephane Belin

## Abstract

Unlike immature neurons and neurons from the peripheral nervous system (PNS), mature neurons from the central nervous system (CNS) cannot regenerate after injury. In the past 15 years, huge progress has been made to identify molecules and pathways necessary for neuroprotection and/or regeneration after CNS injury. In most regenerative models, phosphorylated ribosomal protein S6 (p-RPS6) is upregulated in neurons, which is often associated with an activation of the mTOR pathway. However, the exact contribution of post-translational modifications of this ribosomal protein in CNS regeneration remains elusive. In this study, we demonstrate that RPS6 phosphorylation is essential for PNS and CNS regeneration. We show that this phosphorylation is induced during the preconditioning effect in dorsal root ganglion (DRG) neurons, and that it is controlled by the p90S6 kinase RSK2. Our results reveal that RSK2 controls the preconditioning effect and that the RSK2-RPS6 axis is key for this process, as well as for PNS regeneration. Finally, we demonstrate that RSK2 promotes CNS regeneration in the dorsal column and allows functional recovery. Our data establish the critical role of RPS6 phosphorylation controlled by RSK2 in CNS regeneration and give new insights into the mechanism related to axon growth and circuit formation after traumatic lesion.

## Introduction

Developing neurons or the ones from the peripheral nervous system (PNS) are able to regrow their axon upon lesion. In contrast, mature neurons from the central nervous system (CNS) fail to regenerate their axon after an insult, whether chronic, such as neurodegenerative diseases, or a traumatic (brain and spinal cord injuries) injuries. This leads to irreversible and permanent motor, cognitive and/or sensory disabilities in affected patients. The continuously increasing number of such nervous system disorders worldwide, along with the current absence of efficient therapy, makes neuronal growth and functional recovery major challenges of public health today.

For decades, regeneration failure in the CNS has been explained by the detrimental effect of the environment after injury. Indeed, axon regeneration is impaired by the presence of growth-inhibitory components, such as myelin debris and components of the glia scar. However, their contribution to CNS regenerative failure is limited [1, 2]. In fact, it is now clear that neurons themselves are responsible of this discrepancy. Since 2008, modulation of neuronal intrinsic capacities have unlocked CNS regeneration to some extent [1]. This is the case of the experimental activation of the mTOR pathway via deletion of PTEN (**<u>P</u>**hosphatase and **<u>TEN</u>**sin homologue), which triggers robust axon regeneration in the visual system and in the corticospinal tract [3–7]. Subsequently, combinatorial/synergistic approaches, which often involve mTOR pathway activation have led to long-distance regeneration. For example, the co-activation of mTOR and JAK/STAT pathways, through deletion of PTEN and SOCS3, respectively, allow retina ganglion cells to regenerate from the eye to the optic chiasm after optic nerve injury [8]. Additionally, the study of regenerative capacity of specific subpopulations of RGC revealed osteopontin and IGF control of axon regeneration through mTOR activation [9]. Proteomic analysis of RGC injury response highlighted that several molecular pathways are modulated by the lesion itself, among which the mTOR pathway[10].

One major readout of mTOR activation is the phosphorylation of the ribosomal protein S6 (RPS6) [11], that belongs to the small 40S subunit of the ribosome, the functional unit of protein synthesis in cells. RPS6 is an RNA-binding protein that stabilizes the ribosome by interacting with the ribosomal RNA [12]. Among all ribosomal proteins, RSP6 has attracted most attention as it was the first ribosomal protein shown to have inducible post-translational modifications [13]. Since almost 40 years, the phosphorylation of RSP6 has been studied, yet, many unknowns remain about its physiological functions [14]. Interestingly, in the visual system, RGC subpopulations that are the most resilient to injury have a high level of RPS6 phosphorylation, which is maintained after injury [9]. Nevertheless, whether this phosphorylation is directly associated with mTOR activation in these cells remain unclear. Moreover, in some cases, injury signals may trigger specific events to prime neurons towards a pro-regenerative response. This feature has been elegantly described in the model of dorsal column lesion in the spinal cord [15]. As part of the CNS, the dorsal column, formed by the central branch of dorsal root ganglia (DRG) neurons, is not able to regenerate after spinal cord injury. However, a prior lesion of the DRG peripheral branch, which forms for example the sciatic nerve at the lumbar level, primes DRG neurons to the regeneration of their central branch: this is called the preconditioning effect [16].

The phosphorylation status of RPS6 stands as critical to promote key cellular programs of regeneration. Yet, the exact role of RPS6 phosphorylation and the mechanisms regulating this post-translational modification in the process of CNS regeneration remain elusive. CNS regeneration usually links RPS6 phosphorylation regulation with PTEN deletion, Rheb overexpression or the IGF/Osteopontin signaling, which all converge to mTOR activation [5, 9, 17]. Interestingly, the level of RPS6 phosphorylation increases in DRG neurons after sciatic nerve injury [10, 18], possibly in an mTOR-independent manner [19]. Indeed, mTOR activation is dispensable for the preconditioning effect in DRG neurons, as rapamycin treatment, an inhibitor of mTOR, does not block regeneration [19, 20]. Surprisingly, RPS6 phosphorylation is maintained in these neurons, even upon mTOR inhibition. Furthermore, one target of mTOR, S6 kinase 1 (S6K1), inhibits axon regeneration in *C. elegans* or in mammals through negative feedback on mTOR [21, 22]. Altogether, other pathways controlling RPS6 phosphorylation must be investigated to better understand regeneration mechanisms in the nervous system.

Proteins of the p90 S6 kinase (RSK) family are also known as regulators of RPS6 phosphorylation [23] The RSK protein family is composed of 4 isoforms (RSK1-4), with high homology (from 80% to 87% [24]. RSK are mostly activated by extracellular signal-regulated kinase (ERK) and regulate important processes in cells, such as growth, survival, proliferation and cell cycle progression [24]. However, there is no data available for RSK-RPS6 axis contribution in CNS regeneration. Understanding the precise regulation of RPS6 phosphorylation and the implication of RSK will be key to understand the molecular mechanism underlying CNS regeneration and preconditioning. In this study, we focus on the lumbar DRG as a model of central and peripheral nervous system regeneration. We study a mouse line with unphosphorylable RPS6 to decipher its contribution to regeneration. We show that RPS6 phosphorylation is essential not only for PNS regeneration but also for the preconditioning effect. Among the 4 RSK, RSK2 is strongly expressed by DRG and its expression is regulated by axon injury. We further show that RSK2 modulates RPS6 phosphorylation to promote spinal cord regeneration and functional recovery in mice. Together, our results provide evidence that the RSK/RPS6 is at play in CNS regeneration.

## Results

### RPS6 phosphorylation controls the preconditioning effect and contributes to sciatic nerve regeneration

The phosphorylation of ribosomal protein RPS6 is often used as a readout of mTOR activation [11]. However, the exact contribution of RPS6 during regeneration has never been addressed. Indeed, RPS6 is known as a core ribosomal protein, for which the phosphorylation has been well described [12, 14]. RPS6 has five serine residues that can be phosphorylated (Ser235, Ser236, Ser240, Ser244 and Ser 247) (**Suppl Fig 1A**). To understand the role of RPS6 during axon regeneration, we analyzed its dynamics of phosphorylation upon sciatic nerve injury (**Fig 1A**). We collected the lumbar dorsal root ganglia DRG (L3 to L5) in intact (naive) condition and after 1, 3 and 7 days post-sciatic nerve injury (dpi) in 6 week-old wild-type mice. Western blot analysis using specific anti-p-S6^235-236^ and anti-p-S6^240-244^ antibodies revealed that RSP6 phosphorylation on Ser235-236 is upregulated at 1dpi and reaches a peak at 3dpi, before decreasing at 7dpi (**Fig 1B-C**). On the other hand, RSP6 phosphorylation on Ser240-244 remains overall stable, despite a slight significant increase only at 3dpi (**Fig 1D**). In parallel, we analyzed the levels of phosphorylated RPS6 in DRG using immunofluorescence on cryosections. Consistently with Western blot analysis, we observed an increase of p-S6^235-236^ from 1dpi, with a peak at 3dpi. At 7dpi, Ser235-236 phosphorylation was back to control condition (**Fig 1E-F**). Conversely, the level of p-S6^240-244^ did not display any significant change over time (**Fig 1 G-H**). Together, these results show that RSP6 phosphorylation on Ser235-236 is upregulated in DRG upon sciatic nerve injury.

**Fig 1.**
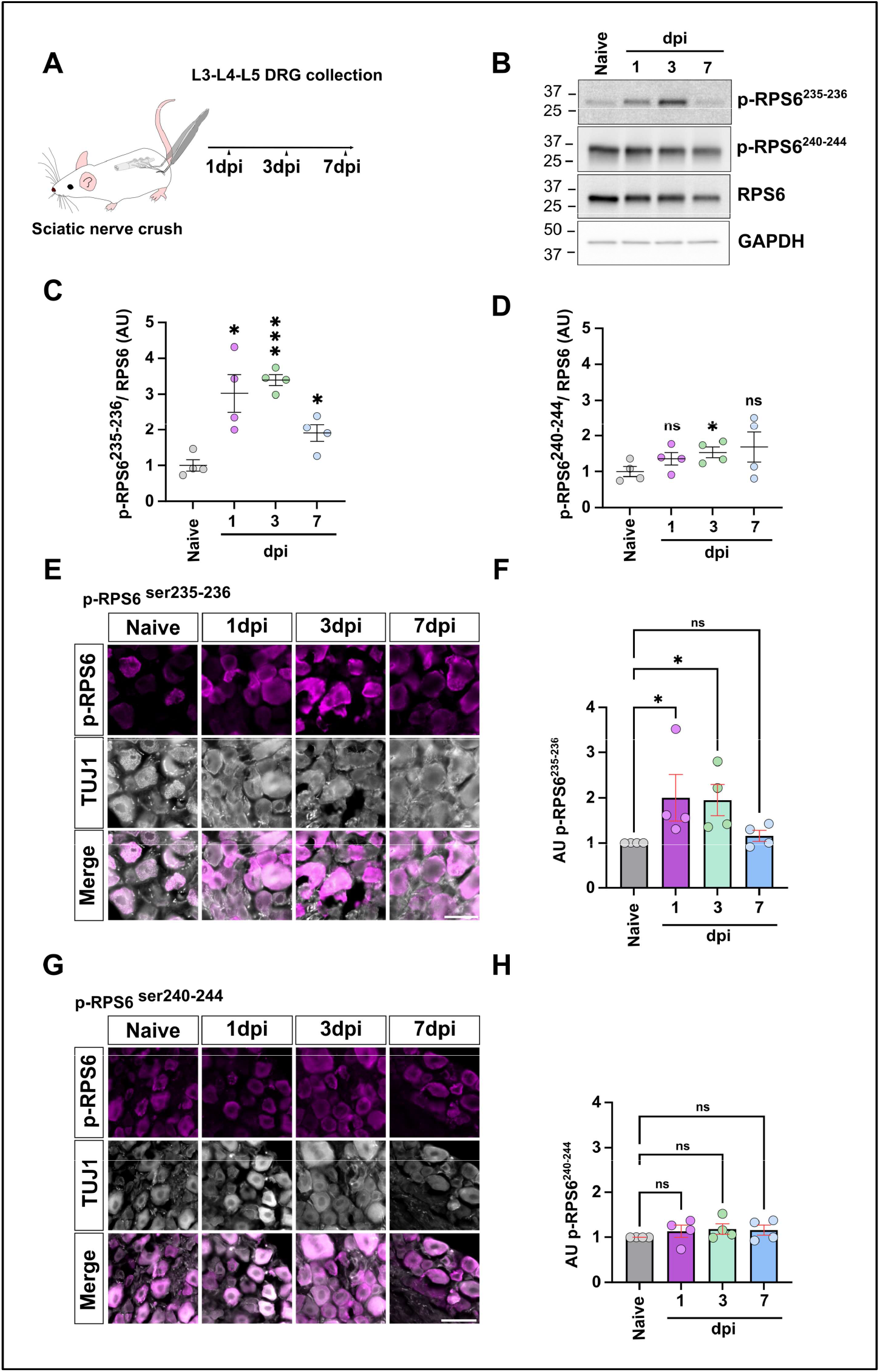
Specific RPS6 phosphorylation on ser235-236 is induced by sciatic nerve injury. (A) Schematic representing the workflow of experiments. (B) Representative western blot showing increase of RSP6 phosphorylation on Ser 235-236 at 1- and 3-days post injury (dpi) while total RSP6 and GAPDH expressions remain stable. (C-D) Graphs showing the quantification of RPS6 phosphorylation on Ser235-S236 (left graph) and on Ser240-244 (right graph) normalized to total RSP6. (mean ± SEM; One-sample t-test; N=4 biological independent animals per group) (E) Representative microphotographs of DRG sections stained with anti-p-RSP6^Ser235-236^ (in magenta) and anti-Tuj1 (in gray) in intact and at different time points upon sciatic nerve injury. Scale bar: 50um. (F) Graphs showing the quantification of E confirming western blot data with an increase of p-RSP6^Ser235-236^ at at 1- and 3-dpi (Mean ± SEM; One-way ANOVA; Dunn’s multiple comparisons test; N=3 biological independent animals per group) (G) Representative microphotographs of DRG sections stained with anti Phospho-RPS6 Ser40-244 (in magenta) and anti-Tuj1 (in gray) in intact and at different time points upon sciatic nerve injury. Scale bar: 50um. (H) Graphs showing the quantification of F (Mean ± SEM; One-way ANOVA; Dunn’s multiple comparisons test; N=3 biological independent animals/group). ⁎⁎⁎ p<0.001, ⁎⁎ p<0.01, ⁎ p<0.05.

In order to assess the contribution of RPS6 phosphorylation in axon regeneration, we analyzed the regenerative response of a transgenic mouse line that carries endogenously an unphosphorylable version of RPS6 [25]. In this mouse line, all serines at 235, 236, 240, 244 and 247 sites are mutated to alanine (**Fig S1A**). We verified that RPS6 cannot be phosphorylated with immunofluorescence on intact DRG (**Fig S1B**). We then extracted proteins from intact and 3dpi DRG in 6 week-old wild-type (RSP6^p+/p+^), heterozygous (RPS6^p+/p-^) and homozygous (RPS6^p-/p-^) littermates. Western blot analysis using p-S6^235-236^ and p-S6^240-244^ antibodies validated that RPS6 is not phosphorylated, neither on 235-236 nor in 240-244 sites, in intact and 3dpi DRG of homozygous mutant mice (RPS6^p-/p-^),in contrast to RSP6^p+/p+^ and RPS6^p+/p-^ littermates (**Fig S1C-E**). The total level of RSP6 was used as a control and did not differ between all genotypes. Consistently with the peaked expression of p-S6^235-236^ at 3dpi (**Fig 1B-C**), we observed a significant increase of Ser235-236 phosphorylation both in RSP6^p+/p+^ and in RPS6^p+/p-^ mice at 3dpi (**Fig S1D**). On the other hand, no change was observed in the level of Ser240-244 phosphorylation at 3dpi (**Fig S1E**).

Next, we asked whether RSP6 phosphorylation was required for the preconditioning effect. To do so, we performed sciatic nerve injury unilaterally on mice from the mutant unphosphorylable RPS6 mouse line. Three days later, we isolated L3 to L5 DRG neurons from the intact (naive) side and injured (preconditioned) side, and cultured them for 16 hours (**Fig 2A**). In naive condition, neurons from RPS6^p+/p+^ and RPS6^p-/p-^ mice have short, highly-ramified neurites in culture. Indeed, we found no significant difference in longest neurite length nor in ramification spacing between RPS6^p+/p+^and RPS6^p-/p-^ naive DRG neurons (**Fig S1F-I**). As expected, in the preconditioned cultures (after sciatic nerve injury), DRG neurons of the RPS6^p+/p+^ genotype have the typical phenotype of preconditioned neurons, with long neurites and few ramifications (**Fig 2B**). Strikingly, in RPS6^p-/p-^ preconditioned DRG neurons, neurites are short and highly ramified, at the same level as in the naive condition (**Fig 2B-E**). In summary, this experiment shows that RPS6 phosphorylation is necessary for the preconditioning effect.

**Fig 2.**
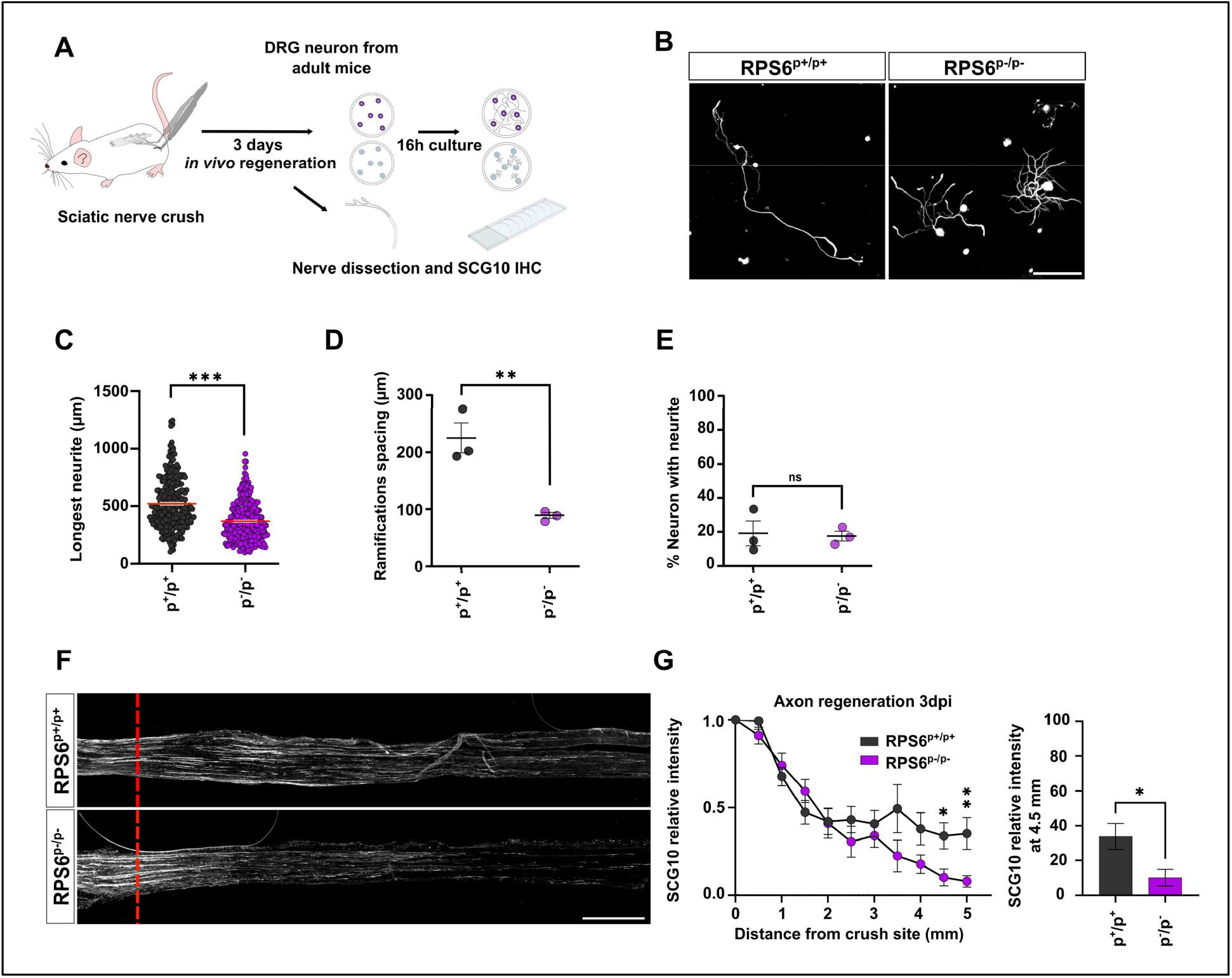
RPS6 phosphorylation is key for axon regeneration. (A) Workflow of unphosphorylable RPS6 mice line analysis in the paradigm of preconditioning and axon regeneration. (B) Representative microphotographs of preconditioned cultured of mature DRG neurons from WT (RPS6^p+/p+^), and homozygous (RPS6^p-/p-^) mice defective for RPS6 phosphorylation showing that in homozygous (RPS6^p-/p-^) mice, the preconditioning effect is inhibited. Scale bar: 250um (C-E) Graphs showing the quantification of B.(C) Longest neurite after per neuron 16h after plating (Mean ± SEM; Unpaired t test with Welch’s correction, 3 independent DRG cultures, approximately 50 cells per conditions per culture). (D) Distance between two ramifications in longest neurite; Mean ± SEM Unpaired t test; 3 independent DRG cultures; approximately 50 cells per conditions per culture d. (E) Percentage of neurons with a neurite 16h after plating Mean ± SEM Unpaired t test; 10 random microscopy fields quantified per conditions per culture.) (D) Representative confocal images of the sciatic nerve sections 3 days post-injury from WT (RPS6^p+/p+^) and homozygous (RPS6^p-/p-^) mice. Regenerating axons are labeled with anti-SCG10 antibody (white). Red dashed line indicates the injury site. Scale bar: 500 mm. (E) Quantification of regenerative axons 3dpi from D (Mean ± SEM; Unpaired Multiple-t-test N=7-8 independent animal per groups). (F) Comparison of regenerative SCG10 axon between RPS6^p+/p+^ and RPS6^p-/p-^. ⁎⁎⁎ p<0.001, ⁎⁎ p<0.01, ⁎ p<0.05.

Finally, we asked whether RPS6 phosphorylation was involved in PNS regeneration. To this end, we performed sciatic nerve injury in 6 week-old RPS6^p+/p+^ and RPS6^p-/p-^ mice and analyzed the extent of regeneration at 3dpi, by SCG10 immunostaining on sciatic nerve sections. Interestingly, while RPS6^p+/p+^ and RPS6^p-/p-^ mice had the same number of axons regenerating in the sciatic nerves, we found that axons from RPS6^p-/p-^ grew significantly less (**Fig 2F-G**). This result suggests that RPS6 phosphorylation is involved in long-distance growth of regenerating PNS axons.

### RPS6 phosphorylation and preconditioning effect are not controlled by the mTOR pathway

RPS6 phosphorylation is commonly used as a readout of mTOR pathway activation, particularly in nervous system regeneration [8, 26]. As RPS6 phosphorylation is key for the preconditioning effect and sciatic nerve regeneration, we asked which signaling pathway controls its phosphorylation in DRG. To do so, we used an *in vitro* pharmacological approach. We performed sciatic nerve crush unilaterally on wild-type mice and three days later, we isolated DRG neurons to plate them. We used cycloheximide as a global inhibitor of translation, rapamycin and Torin 1 as inhibitors of mTOR, PF-4708671 as a S6 kinase inhibitor and BRD7389 as an inhibitor of the p90 RSK (**Fig 3A**) and used DMSO as control [27]. One hour after plating, we treated cultures with the drug of interest, then we assessed neuronal growth after 16 hours. We found that cycloheximide-mediated inhibition of global translation totally blocks axon outgrowth, both in naive and in preconditioned DRG cultures (**Fig S2A-C and Fig 3B-C**). This result shows that protein translation is key for axon outgrowth in naive and preconditioned cultures.

**Fig 3.**
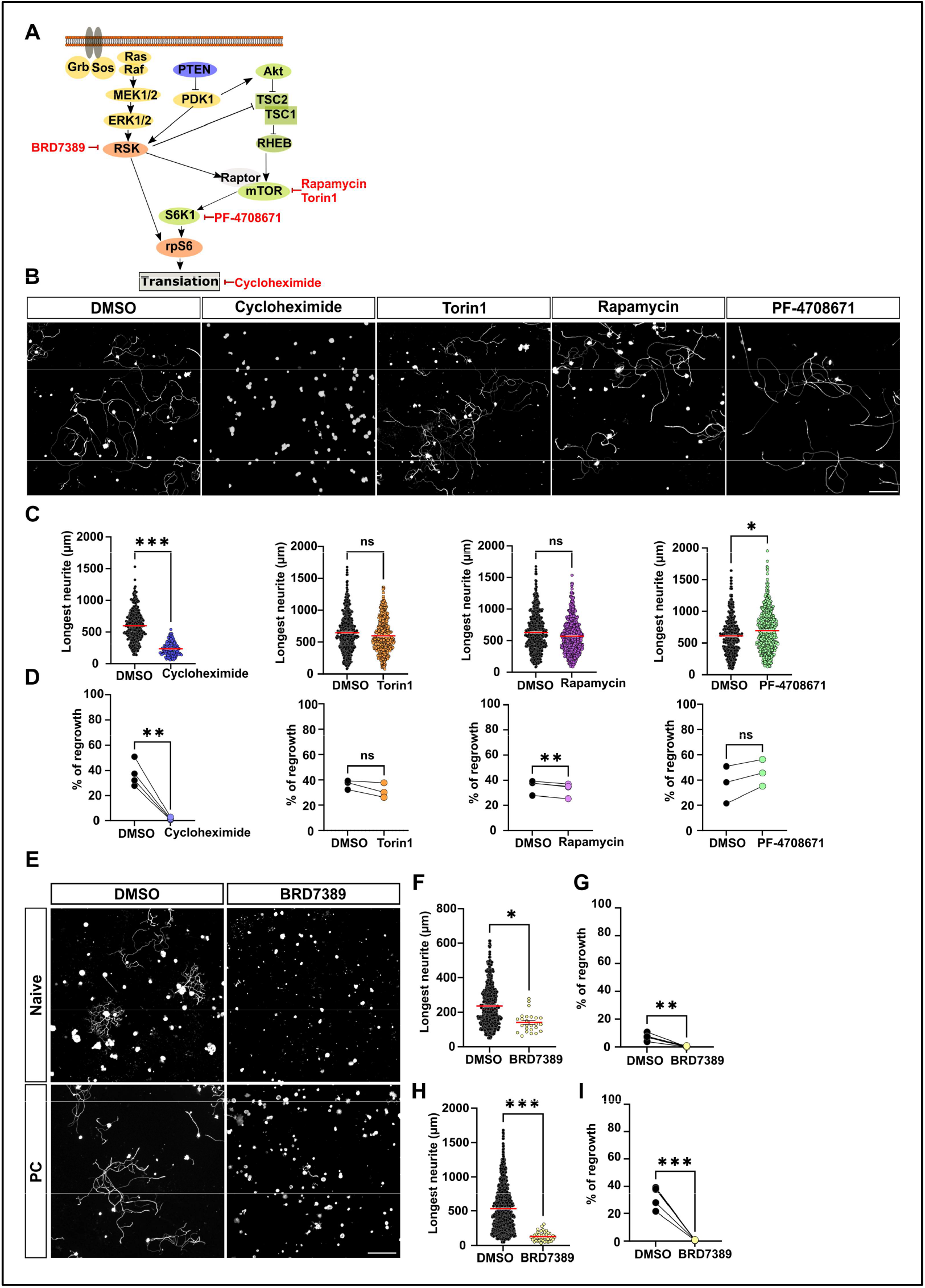
RSK controls the preconditioning effect in mature DRG neurons. (A) Ribosomal S6 kinase simplified pathway and inhibitors used in this study (in red) (B) Representative microphotographs of preconditioned cultures of mature DRG neurons treated with DMSO (control), transcription inhibitor (cycloheximide-2nM), mTOR inhibitors (Torin-5nM) or Rapamycin-0.1nM), S6K1 inhibitor (PF-4708671-8uM) Scale bar: 250 um. (C) Longest neurite quantification per neuron 16h after plating (Mean ± SEM; Kruskal wallis test Dunn’s multiple comparisons test; 3-4 independent DRG cultures; approximately 50-100 cells per condition per culture (except for cyclohexamide). (D) Percentage of neurons with a neurite 16h after plating Mean ± SEM Paired t test; 10 random microscopy fields quantified per conditions). (E) Representative microphotographs of naive and preconditioned cultures of mature DRG neurons treated with DMSO (control) or RSK inhibitor (BRD-7389 (3uM)). Scale bar: 250um. (F) Quantification of the longest neurite per neuron 16h after plating in naive condition. (Mean ± SEM Kruskal Wallis test Dunn’s multiple comparisons test, 5 independent DRG cultures; approximately 50-100 cells per condition per culture for DMSO condition; all neurons found with a neurite were quantified in BRD7389 condition) (G) Percentage of neurons with a neurite 16h after plating in naive DRG treated with DMSO or BRD7389. (Mean ± SEM Paired t test; 10 random microscopy fields quantified per condition) (H) Quantification of the longest neurite 16h after plating in PC DRG treated with DMSO or BRD7389 (Mean ± SEM; Unpaired t test with Welch’s correction approximately 50-100 cells per condition per culture for DMSO condition; all neurons found with a neurite were quantified in BRD7389 condition) (I). Percentage of neurons with a neurite 16h after plating in naive DRG treated with DMSO or BRD7389. (Mean ± SEM Paired t test; 10 random microscopy fields were quantified per condition; 5 independent DRG cultures.⁎⁎⁎ p<0.001, ⁎⁎ p<0.01, ⁎ p<0.05.

Interestingly, inhibition of mTOR or S1 kinase did not prevent neurite outgrowth in naive conditions. We found no difference in the length of the longest neurite, nor in the total number of neurons that grow a neurite between control and mTOR inhibition (Torin 1, Rapamycin) treatments. Inhibition of S6K with PF-4708671 caused a slight increase of the number of neurons that grow a neurite (9.0±1.2% for PF-4708671 vs 6.7±1.6% for DMSO, Paired t-test, p=0.0420). In preconditioned DRG cultures, mTOR inhibition had no effect on the extent of growth and ramification of growing neurons. When preconditioned neurons are treated with S6K inhibitor, we observe a slight increase of the longest neurite length (694um ± 13 for PF-4708671 vs 610um ± 15 for DMSO, Mean ± SEM One-way ANOVA, p=0.0196) but the total number of neurons growing a neurite was unchanged (**Fig 3B-D**). Altogether our results show that neither mTOR nor its downstream effector S6K1 are required for the preconditioning effect.

Strikingly on the other hand, inhibition of the RSK family with BRD7389 completely blocked neurite outgrowth, both in naive DRG cultures and in preconditioned DRG cultures (**Fig 3E-I**). We verified that this effect was not due to toxicity by controlling that the number of Tuj1-positive cells is similar between DMSO and BRD7389 treatments. This result suggests that RSK is the family of kinases involved in the preconditioning effect.

### RSK2 expression is regulated by sciatic nerve injury and controls RPS6 phosphorylation in DRG neurons

As BRD7389 treatment shows a striking effect on neuronal growth, we next assessed the expression of RSK family members in adult DRG. In mice, RSK family is composed of 4 isoforms with high homology, particularly in the two kinase domains (**Fig S3A-B**). Therefore, we designed specific RNA probes that target unique and specific regions of each isoform (RSK1 to 4) (**Fig S3C and Table S1**). We performed in situ hybridization on cryosections of adult DRG from 6 week-old wild-type mice (**Fig S3D**). We found that RSK 2 and 3 are enriched in DRG in intact conditions, whereas RSK1 and 4 are lowly expressed (**Fig S3E**).

Then we investigated whether the expression of RSK2 and 3 are modulated by axon injury. To do so, we collected DRG at different time points after sciatic nerve crush (**Fig S3D**). We found that RSK2 is specifically regulated by sciatic nerve injury. Indeed, using in situ hybridization, we observed that *RSK2* mRNA is upregulated at 1dpi and 3dpi, before going back to control (intact) level at 7dpi. In contrast, *RSK3* mRNA expression is not modulated by the injury (**Fig S3F**). Therefore, we focused our study on RSK2.

We then sought to determine the dynamics of RSK2 protein expression in DRG upon sciatic nerve injury (**Fig 4B-E**). We found a significant increase of RSK2 expression from 1dpi, that reaches a peak at 3dpi and then decreases at 7dpi back to control (intact) level, both using Western blot analysis (**Fig 4B-C**) and immunostaining (**Fig 4D-E**) using an anti-RSK2 specific antibody. Importantly, RSK2 dynamics of expression (**Fig 4B-E**) matches RPS6 dynamics of phosphorylation upon sciatic nerve injury (**Fig 1B-C and 1E-F**). This result supports the hypothesis that RSK2 is involved in RPS6 phosphorylation and in the control of the preconditioning effect.

**Fig 4.**
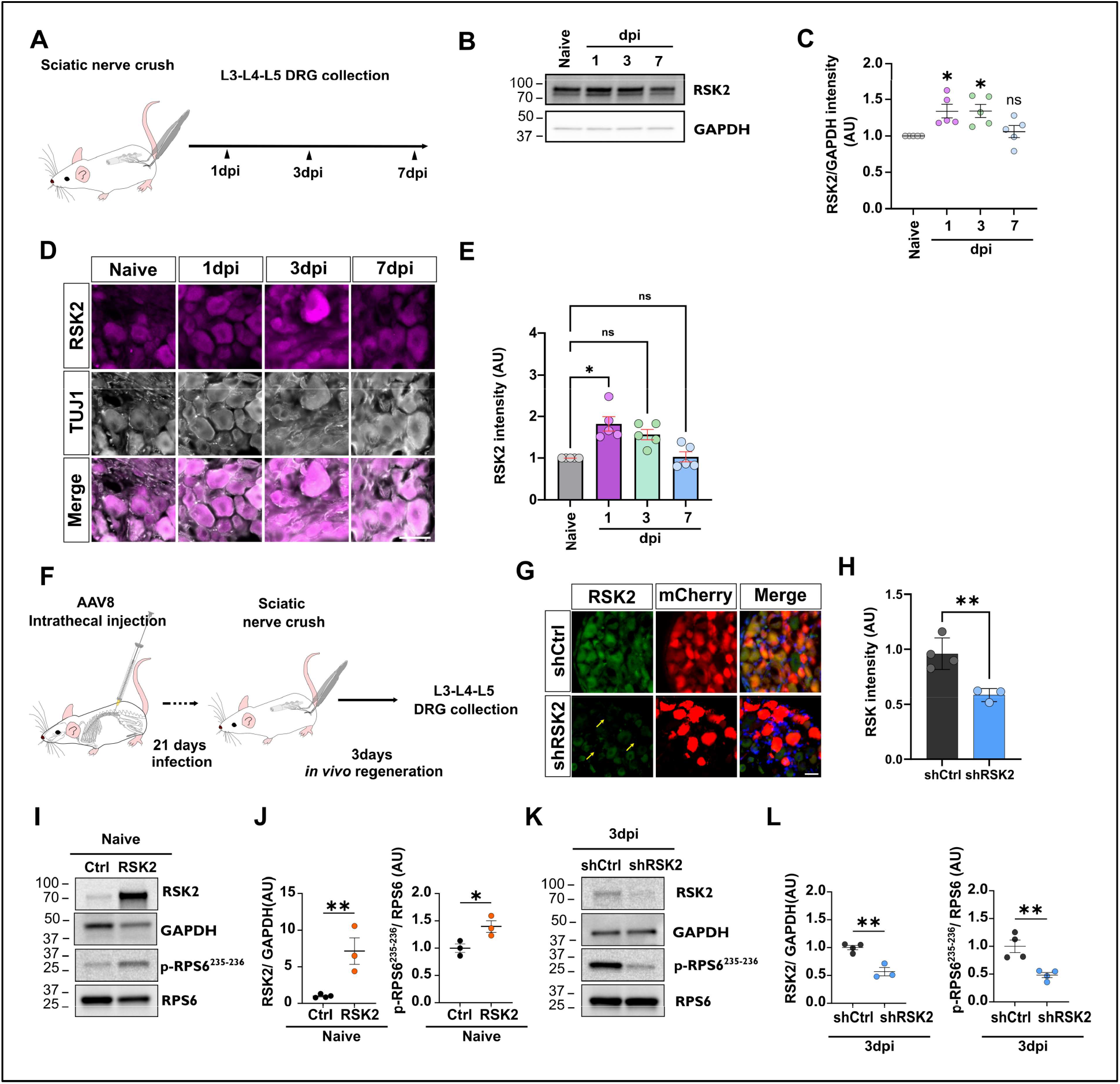
RSK2 regulates RPS6 Ser235-236 phosphorylation in mature DRG. (A)Workflow of experiment. (B) Western blot showing upregulation of RSK2 expression. (C) Quantification of B (Mean ± SEM One sample t-test; N=5 biological independent animal/groups). (D) Representative images of DRG section from intact or 1dpi, 3dpi and 7dpi stained with anti-RSK2 (in magenta) and anti-Tuj1 (in gray), Scale bar:50 um. (E) Quantification of D. (One-way ANOVA; Dunn’s multiple comparisons test; N=5 individual animals per group) (F) Timeline of the experimental procedure to investigate *in vivo* role of RSK2 in RPS6 phosphorylation. (G) Representative images of infected DRG by shRSK2 or shCtrl stained with anti-RSK2 (in green) and anti-RFP (in red); Scale bar: 50um. (H) Quantification of G. (Mean ± SEM; Unpaired t-test; N=3-4 individual animals per group). (I) Western blot showing that overexpression of RSK2, *in vivo* in naive DRG, induces RPS6 phosphorylation on Ser235-236 without sciatic nerve injury. (H) Quantification of G, (Mean ± SEM; One sample t test N=3-4 individual animals per group) (I) Western blot showing overexpression of RSK2 *in vivo* in naive DRG. (J) Quantification of I. (Mean ± SEM; One sample t test N=3-4 individual animal/groups). (K) Western blot showing inhibition of RSK2 *in vivo* in preconditioned DRG 3 days after sciatic nerve injury. (L) Quantification of L; (Mean ± SEM; One sample t test N=3-4 individual animal per groups). ⁎⁎⁎ p<0.001, ⁎⁎ p<0.01, ⁎ p<0.05.

In order to study the regulation of RPS6 phosphorylation by RSK2, we generated AAV viral vectors that (i) on one hand, overexpresses RSK2 and (ii) on the other hand, inhibits specifically RSK2 expression with an shRNA-based silencing approach (shRSK2) (**Fig S4A**). We modulated RSK2 expression in DRG *in vivo* by using intrathecal injection of AAV8-shRSK2 in 4 week-old wild-type mice (**Fig 4F**). *In vivo* overexpression of RSK2 in DRG significantly enhanced RPS6 phosphorylation at Ser235-236 in naive condition (**Fig 4I**), to the same level of RPS6 phosphorylation observed at 3dpi (**Fig 1**).

For the silencing approach, we first verified shRSK2 specificity by co-transfecting it with plasmids overexpressing RSK1, RSK2, RSK3 or RSK4 in N2A cells (**Fig S4A**). Western blot analysis confirmed that shRSK2 inhibits RSK2 expression only (**Fig S4B-E**). Three weeks after intrathecal injection of AAV8-shRSK2, more than 90% of DRG were infected with the virus (**Fig 4G**) and RSK2 expression was decreased by 50% (**Fig 4H**). Strikingly, RSK2 inhibition blocked the phosphorylation of RPS6 at Ser235-236 normally induced by sciatic nerve injury (**Fig 4K-L**). Together, our results highlight RSK2 as the main kinase that controls RPS6 phosphorylation in DRG upon sciatic nerve injury.

### RSK2 controls the preconditioning effect and sciatic nerve regeneration

Next, we asked whether RSK2 was involved in the preconditioning effect. To this end, we modulated RSK2 expression in vivo by intrathecal injection of AAV8 vector, and analyzed the regeneration potential of both naïve and preconditioned DRG in culture (**Fig S4A**). Strikingly, overexpression of RSK2 *in vivo* caused naive DRG to grow significantly longer neurites with significantly fewer ramifications than naive control, a phenotype that is identical to the preconditioning effect (**Fig 5A-D**). We found that this effect is specific of RSK2, as overexpression of RSK3 in naive DRG does not mimic the preconditioning effect (**Fig S5A-E**). Conversely, inhibition of RSK2 expression in vivo resulted in the loss of the preconditioning effect in DRG of the sciatic nerve injured side. Indeed, in absence of RSK2, preconditioned DRG neurons resemble naive ones, with shorter, highly-ramified neurites (**Fig 5G-J).**

**Fig. 5.**
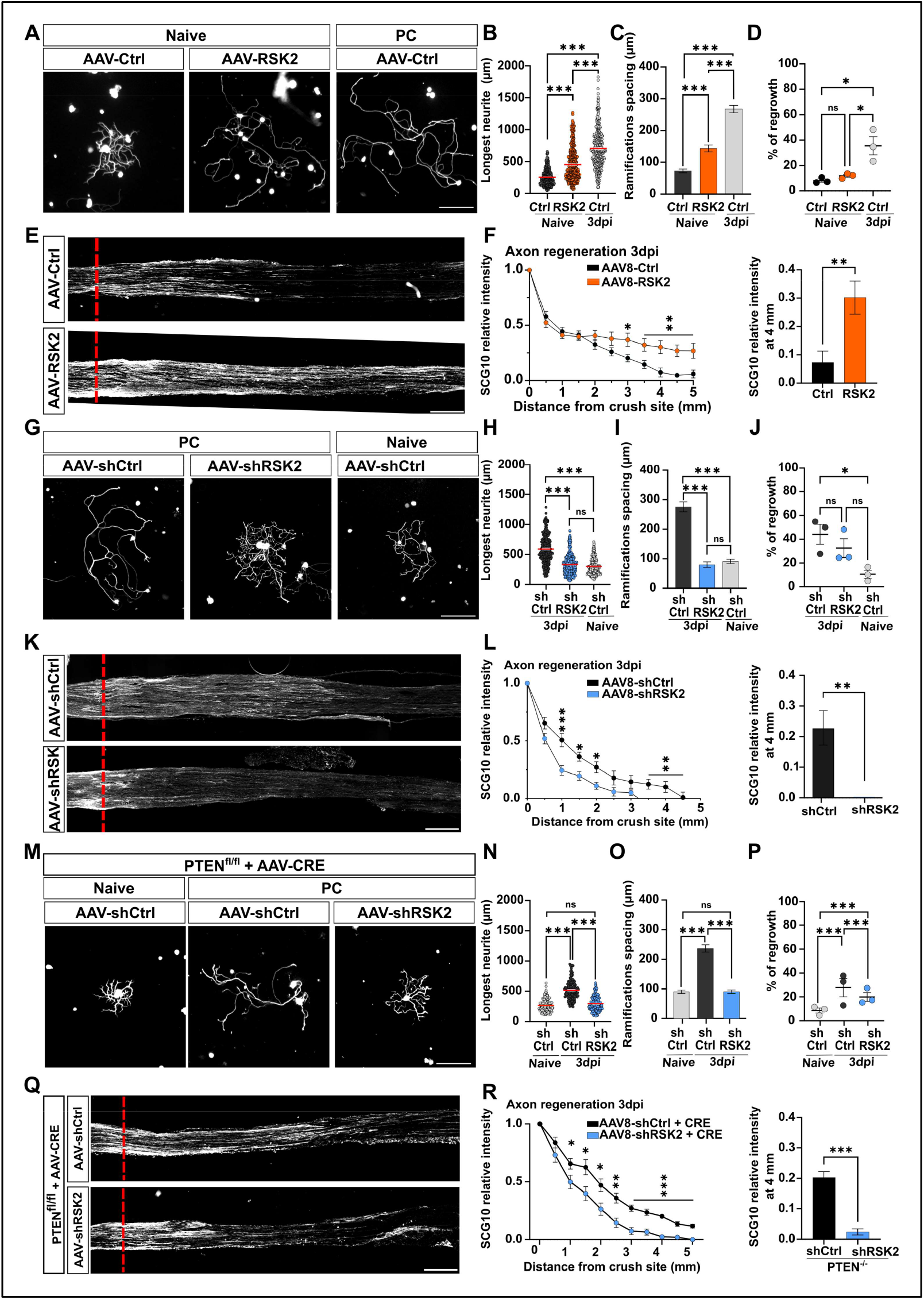
RSK2 controls the preconditioning effect and axon regeneration in the PNS. (A) Representative microphotographs of DRG dissociated cultures showing that RSK2 overexpression in naive cultures phenocopies the preconditioning effect. Scale bar:250um. (B-D) Quantification of A: (B) Longest neurite per neuron 16h after plating (Mean ± SEM; One-way ANOVA Dunn’s multiple comparisons test, 3 independent DRG cultures, approximately 50-100 cells counted per condition per culture). (C) Mean distance between two ramifications (Mean ± SEM Two-way ANOVA; Tukey’s multiple comparisons test; 3 independent DRG cultures; approximately 50 cells per condition per culture) and (D) Percentage of neurons with a neurite 16h after plating (Mean ± SEM Two-way ANOVA Tukey’s multiple comparisons test; 10 random microscopy fields quantified per condition per culture). (E) Representative images of the sciatic nerve sections 3 days post-injury from mice intrathecally injected with AAV8-PLAP (control) or AAV8-RSK2. Regenerating axons are labeled with anti-SCG10 antibody (white). Red dashed line indicates the injury site. Scale bar: 500 um. (F) Quantification of regenerative axons from E. (Mean ± SEM; Multiple t-test, at least 6 individual animal per condition). (G) Representative microphotograph of DRG dissociated cultures showing that RSK2 inhibition in preconditioned cultures phenocopies the naive condition. Scale bar:250um (H-J) Quantification of G. (H) Longest neurite per neuron 16h after plating (Mean ± SEM One-way ANOVA Dunn’s multiple comparisons test, 3 independent DRG cultures, approximately 50-100 cells count per condition per culture). (I) Mean distance between two ramifications (Mean ± SEM Two-way ANOVA Tukey’s multiple comparisons test; 3 independent DRG cultures; approximately 50 cells per condition per culture) and (J) Percentage of neurons with a neurite 16h after plating (Mean ± SEM Two-way ANOVA Tukey’s multiple comparisons test; 10 random microscopy fields were quantified per conditions). (G) Representative images of the sciatic nerve sections 3 days post-injury from mice injected intrathecally with AAV8-Sh-Scrambled or AAV8-Sh-RSK2. Regenerating axons are labeled with anti-SCG10 antibody (white). Red dashed line indicates the injury site. (H) Quantification of regenerative axons from G. (Mean ± SEM Multiple t-test, at least 6 individual animals per condition). (M) Representative microphotographs of DRG dissociated cultures showing that RSK2 inhibition in PTEN deleted preconditioned cultures phenocopies the naive condition. Scale bar: 250um (N-P) Quantification of M. (N) Longest neurite per neuron 16h after plating (Mean ± SEM One-way ANOVA Dunn’s multiple comparisons test, 3 independent DRG cultures, approximately 50-100 cells counted per conditions per culture); (O) Mean distance between two ramifications (Mean ± SEM Two-way ANOVA Tukey’s multiple comparisons test; 3 independent DRG cultures; approximately 50 counted cells per conditions per culture) and (P) Percentage of neurons with a neurite 16h after plating (Mean ± SEM Two-way ANOVA Tukey’s multiple comparisons test; 10 random microscopy fields were quantified per condition). (Q) Representative images of the sciatic nerve sections 3 days post-injury from mice injected intrathecally with AAV8-Sh-Scrambled or AAV8-Sh-RSK2 and AAV8-CRE in PTENfl/fl mice. Regenerating axons are labeled with anti-SCG10 antibody (white). Red dashed line indicates the injury site. (R) Quantification of regenerative axons from K. (Mean ± SEM Multiple t-test, at least 6 individual animal per condition).⁎⁎⁎ p<0.001, ⁎⁎ p<0.01, ⁎ p<0.05.

In parallel, we analyzed the regeneration potential of DRG in the sciatic nerve *in vivo*. We injected intrathecally AAV8-RSK2, AAV-ShRNA-RSK2 or corresponding controls in 4 week-old animals, and performed unilateral sciatic nerve crush two weeks later **(Fig S5A)**. The extent of axon regeneration was analyzed by SCG10 immunostaining at 3dpi. Similarly to the effect on DRG cultures, we found that RSK2 overexpression enhances sciatic nerve regeneration (**Fig 5E-F**), with axons extending up to 5mm from the lesion site. This effect is specific of RSK2, as overexpression of RSK3 did not affect sciatic nerve regeneration (Fig S5D-E). In contrast, RSK2 inhibition blocks axon regeneration in the sciatic nerve (**Fig 5K-L**). Together, our results demonstrate that RSK2 is critical for peripheral nerve regeneration.

To rule out the contribution of mTOR in the preconditioning effect, we activated the mTOR pathway through intrathecal injection of AAV8-Cre in PTEN^f/f^ mice. Deletion of PTEN leads to activation of the mTOR pathway [26]. As expected, mTOR activation in naive DRG neurons does not induce the preconditioning effect. We then analyzed the growth outcome of RSK2 inhibition together with mTOR activation in preconditioned DRG neurons. We observed that mTOR activation does not modify the preconditioned effect. In contrast, inhibition of RSK2 in PTEN-deleted neurons blocks the preconditioning effect: neurons grow shorter and highly ramified neurites, similarly to the naive condition (**Fig 5M-P**). In parallel, we analyzed axon regeneration of sciatic nerve in these mice. We found that mTOR pathway activation does not counteract the suppression of axon regeneration induced by RSK2 inhibition (**Fig 5Q-R**).

Altogether, our results show that RSK2 controls the preconditioning effect and PNS regeneration independently of mTOR.

### RSK2 controls the preconditioning effect via RPS6 phosphorylation

Our results show that RSK2 regulates RPS6 phosphorylation. Moreover, RSK2 and p-RPS6 are both indispensable for the preconditioning effect (**Fig 2 and Fig 5**). Thus, we asked whether RSK2 regulates the preconditioning effect via RPS6 phosphorylation. To do so, we overexpressed RSK2 in DRG of the unphosphorylable RPS6 mutant mouse line (**Fig S1A**), by injecting intrathecally AAV8-RSK2 or control in RPS6^p+/p+^ and RPS6^p-/p-^ 4 week-old mice. Two weeks later, we performed unilateral sciatic nerve injury. We first analyzed DRG cultures at 3dpi. As expected, in naive RPS6^p+/p+^ DRG condition, RSK2 overexpression induces a preconditioning effect-like phenotype (**Fig 6A-B**). Conversely, RSK2 overexpression in naive RPS6^p-/p-^ DRG does not produce the preconditioning effect: neurites are short and highly ramified, as naive control culture (**Fig 6A-D**). Very interestingly, in preconditioned RPS6^p-/p-^ DRG cultures, RSK2 overexpression has no effect on neurons as they maintain their naive phenotype (**Fig 6A-D**). Our results show that RSK2-mediated control of the preconditioning effect depends on RPS6 phosphorylation. In parallel, analysis of regeneration in the injured sciatic nerve showed that, in contrast to RPS6^p+/p+^ mice, overexpression of RSK2 in RPS6^p-/p-^ mice completely switches off axon regeneration (**Fig 6E-F**). Altogether, our results show that the RSK2-RPS6 axis is critical for the preconditioning effect and peripheral nervous system regeneration.

**Fig 6.**
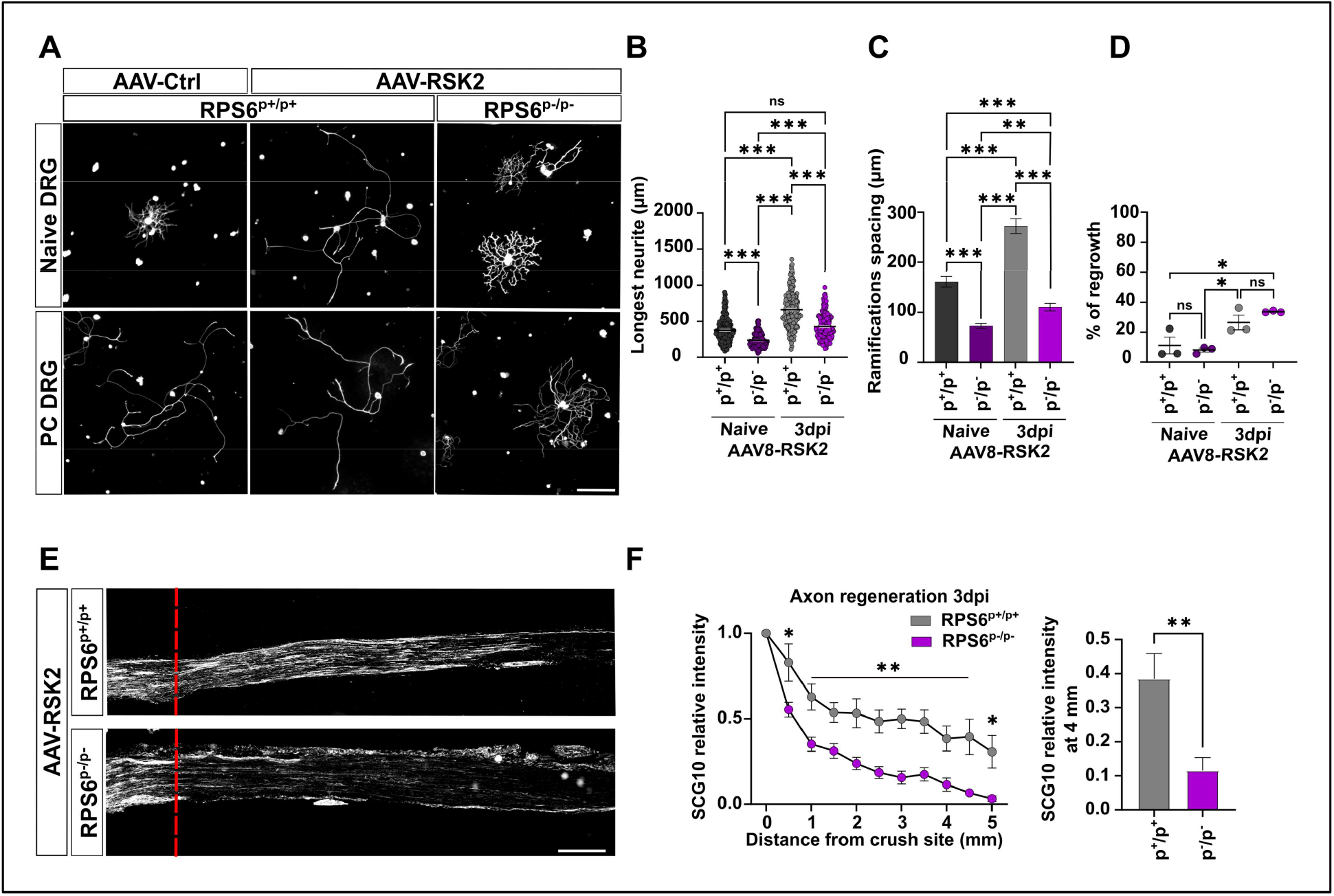
RSK2 needs a phosphorylatable RPS6 to induce the preconditioning effect and axon regeneration in the PNS. (A) Representative microphotographs of DRG dissociated cultures showing that RSK2 overexpression phenocopies the preconditioning effect in naïve cultures from RPS6^p+/p+^ mice but not from RPS6^p-/p-^ cultures; Scale bar: 250um. (B-D) Quantification of A: (B) Longest neurite per neuron 16h after plating (Mean ± SEM One-way ANOVA; Dunn’s multiple comparisons test, 3 independent DRG cultures; approximately 50 counted cells per condition per culture); (C) Mean distance between two ramifications (Mean ± SEM One-way ANOVA; Dunn’s multiple comparisons test, 3 independent DRG cultures; approximately 50 cells counted per condition per culture) (D) Percentage of neurons with a neurite 16h after plating (Mean ± SEM; One-way ANOVA Dunn’s Multiple comparison test; 10 random microscopy fields were quantified per condition). (E) Representative confocal images of the sciatic nerve sections 3 days post-injury from RPS6^p+/p+^ or RPS6^p-/p-^ mice injected intrathecally with AAV8-RSK2. Regenerating axons are labeled with anti-SCG10 antibody (white). Red dashed line indicates the injury site. Scale bar: 500 mm. (F) Quantification of regenerative axons from C (Mean ± SEM Multiple t-test, at least 3 individual animals per condition).⁎⁎⁎ p<0.001, ⁎⁎ p<0.01, ⁎ p<0.05.

### RSK2 controls spinal cord regeneration and functional recovery

As the RSK2/RPS6 axis controls the preconditioning effect, we then asked whether it also controls CNS regeneration. To address this question, we focused on the sensory axons that form the dorsal column of the spinal cord. This bundle contains the central branch of the DRG. Like all CNS axons, these axons are unable to regenerate spontaneously after spinal cord injury [15, 16]. Interestingly, the prior lesion of the sciatic nerve (the preconditioning paradigm) promotes axon regeneration in the spinal cord [15]. In this context, we first assessed the dorsal column regeneration in the unphosphorylable RPS6 mutant mouse line. To answer this question, 6 week-old RPS6^p+/p+^ and RPS6^p-/p-^ mice received unilateral sciatic nerve injury and one day after, dorsal column crush injury. One week before sacrifice, we injected Alex555-conjugated cholera toxin B (CTB) into the sciatic nerve, in order to assess dorsal column regeneration Only animals with complete lesions were analyzed, as verified at the cervical level (**Fig S6A-B**). Regeneration was analyzed 6 weeks after spinal cord injury. As expected, in RPS6^p+/p+^ mice, the preconditioning lesion of the sciatic nerve induces regeneration in the dorsal column. In RPS6^p-/p-^ mice, on the other hand, this regenerative effect is completely abolished (**Fig S6C**). This further confirms that RPS6 phosphorylation is essential to trigger axon regeneration in the dorsal column.

Since RSK2 controls the preconditioning effect via RPS6 phosphorylation, we asked whether RSK2 overexpression is sufficient to induce axon regeneration in the spinal cord. Thus, we injected intrathecally AAV8-RSK2 or AAV8-control in 4 week-old wild-type mice. Two weeks later, we performed crush injury of the dorsal column. (**Fig 7A**). For each sample, analysis of cervical sections confirmed that the lesion was complete (**Fig S6**). In control condition, axons reached the border of the lesion, with few axons observed within the injury site. No axon was found crossing the injury site (**Fig 7B-C**). When RSK2 is overexpressed in DRG (without the preconditioning paradigm), not only do axons enter the lesion, they also cross it and grow beyond the injury site. This phenotype is observed at 6 weeks post-injury and is exacerbated at 8 weeks post-injury (**Fig 7B-E**).

**Figure 7:**
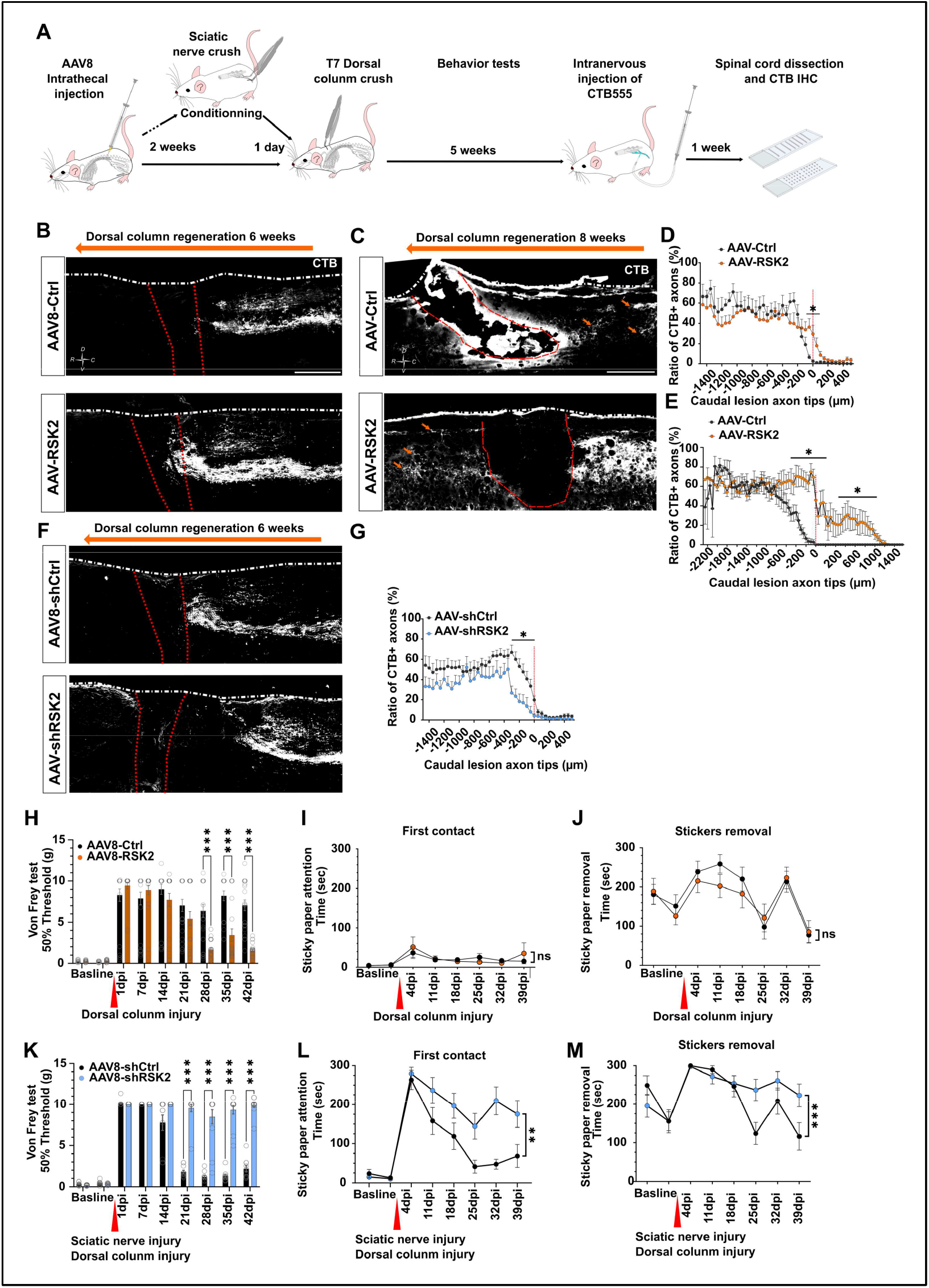
RSK2 induces dorsal column regeneration with functional sensory recovery. (A) Workflow of experiments. (B) Representative confocal images of thoracic spinal cord sagittal sections 6 weeks after dorsal column crush from mice injected intrathecally with AAV8-PLAP or AAV8-RSK2. Regenerative axons are labeled with anti-CTB antibody (white). (C) Representative confocal images of thoracic spinal cord sagittal section 8 weeks after dorsal column crush from mice injected intrathecally with AAV8-PLAP or AAV8-RSK2. Regenerative axons are labeled with anti-CTB antibody (white). (D) Quantification of axon regeneration and dieback from caudal marge of crush site from B. (Mean ± SEM; Mann-Whitney test; N= at least 6 independent animals per condition. (E) Quantification of axon regeneration and dieback from caudal marge of crush site from C. (Mean ± SEM; Mann-Whitney test; N= at least 4 independent animals per condition. (F) Representative confocal images of thoracic spinal cord sagittal section 6 weeks after sciatic nerve crush and dorsal column crush from mice injected intrathecally with AAV8-Sh-Scrambled or AAV8-Sh-RSK2. Regenerative axons are labeled with anti-CTB antibody (white). (G) Quantification of axon regeneration and dieback from caudal marge of crush site from F. (Mean ± SEM; Mann-Whitney test; N= At least 8 independent animals per group). (H) Von Frey experiment to test nociception in mice intrathecally injected with AAV8-PLAP or AAV8-RSK2 2 weeks before and 6 weeks after dorsal column crush; stimulus intensity is showed in grams. (Mean ± SEM; Multiple t test. *N*= At least 11 biologically independent animals per group, each paw was considered independently). (I-J) Sticky paper attention test (first contact and removal) in mice intrathecally injected with AAV8-PLAP or AAV8-RSK2 2 weeks before and 6 weeks after dorsal column crush. (Mean ± SEM; Two-way ANOVA; *N* = At least 11 biologically independent animals per group; NS= non-significant). (K) Von Frey experiment to test nociception in mice intrathecally injected with AAV8-sh-Scramble or AAV8-sh-RSK2 2 weeks before and 6 weeks after sciatic nerve injury and dorsal column crush, stimulus intensity is showed in grams; (Mean ± SEM; multiple t test at least 11-12 biologically independent animals per group, only the injured paw was considered). (L-M) Sticky paper attention test (first contact and removal) in mice intrathecally injected with AAV8-sh-Scrambled or AAV8-sh-RSK2, 2 weeks before and 6 weeks after left sciatic nerve injury and dorsal column crush. (Mean ± SEM Two-way ANOVA at least 11-biologically independent animals per group), ⁎⁎⁎ p<0.001, ⁎⁎ p<0.01, ⁎ p<0.05.

We next assessed whether this regeneration can sustain functional recovery. To this end, we performed two behavioral assays to study sensitive function recovery: the sticky paper attention test (where first contact and total time for removal sticky paper was measured) and the Von Frey filament test. In the sticky paper test, we did not see any difference between control and RSK2 overexpression groups (**Fig 7I-J**). Interestingly, the Von Frey test revealed that mice overexpressing RSK2 have better functional recovery. Indeed, immediately after dorsal column injury, we observed a loss of sensitivity in both groups. While this loss of sensory function was maintained in the control group throughout the whole experiment, the RSK2 overexpression group recovered sensitivity from 28 days after injury (**Fig 7H**). Together, our histological and behavioral analyses show the RSK2 upregulation induces CNS axon regeneration and functional recovery.

To confirm these findings, we tested the effect of RSK2 inhibition on CNS regeneration after preconditioning. In this experiment, 4 week-old wild-type animals received an intrathecal injection of AAV8-shRSK2 or control. Two weeks later, we performed unilateral sciatic nerve injury and the next day, we performed dorsal column crush injury. One week before sacrifice, we injected Alex555-conjugated CTB -– into the sciatic nerve, upstream to the injury site (medial to the spinal cord), in order to assess dorsal column regeneration (**Fig 7A**). For each sample, analysis of cervical sections confirmed that the lesion was complete (**Fig S6**). In control mice, regenerating axons reach the lesion site and some axons are able to cross it, as previously reported [15] (**Fig 7F-G**). When RSK2 is inhibited in DRG, despite the preconditioning paradigm, we observed a massive retraction of the axon bundle from the lesion site. No axon could reach the injury site (**Fig 7F-G**). In order to assess the effect of RSK2 inhibition on sensory functional recovery, we performed the same behavioral tests as described above. In both the Von Frey test and the sticky removal test, inhibition of RSK2 significantly impairs functional recovery induced by the preconditioning (**Fig 7K-M**).

Altogether, our results demonstrate that the RSK2/RSP6 axis is required for sensory axon regeneration in the spinal cord.

## Discussion

Because of current lack of efficient therapies for CNS injury, CNS regeneration represents a major challenge in the field of neurobiology today. Despite major advances in the modulation of intrinsic regrowth in the past 15 years [1, 8, 10, 17], functional recovery has not been unlocked yet. A key pathway that was shown to trigger CNS regeneration, both in the injured visual system and in the injured corticospinal tract, is the mTOR pathway [4, 26]. However, not only the precise mechanisms by which mTOR leads to axon regeneration remain elusive, but the exact contribution of one of its major effectors, phosphorylated RPS6, is unknown.

In our study, we demonstrate that RPS6 phosphorylation is critical for the regeneration of both the PNS and the CNS. Indeed, we show that RPS6 phosphorylation is induced after sciatic nerve injury and is required for the preconditioning effect observed in DRG neurons. In addition, we demonstrate that this phosphorylation is not controlled by mTOR but by the p90S6 kinase RSK2. Members of the RSK family are the first kinases described to phosphorylate RPS6 at the position Ser235-236 [13]. Our results show that the RSK2/RPS6 phosphorylation axis controls the process of preconditioning and PNS regeneration. Moreover, RSK2 promotes regeneration of the central branch of DRG axons in the spinal cord, and associated functional recovery. Altogether, our work sheds light on the critical role of RPS6 phosphorylation and on the importance of its post-translational regulation by RSK2. In this study, we aimed to understand the contribution of RSP6 phosphorylation to the nervous system regeneration. Increased RSP6 phosphorylation correlates with enhanced regenerative capacity, both in the CNS and the PNS [18, 26]. In the CNS, high levels of endogenous RPS6 phosphorylation in specific RGC subpopulations (eg alpha-RGC, ipRGC) correlate with better resilience and regenerative potential after injury [9, 28]. Other neurons like DRG neurons express phosphorylated RPS6. This level increases upon sciatic nerve injury. Moreover, the level of RPS6 phosphorylation decreases in neurons during development and ageing, which again correlates with the decrease of regenerative ability [26, 29]. All these observations suggest a strong connection between the phosphorylation state of RPS6 and the regenerative outcome. Yet, very few studies have addressed the role of RPS6 post-translational modifications - specifically phosphorylation - in axon regeneration.

RPS6 is a ribosomal protein (RP) that belongs to the 40S subunit of the ribosome. It is one of the best studied RPs. Indeed RPS6 is the first RP for which inducible post-translational modification has been reported upon liver injury [13]. The role of RPs during regulation of protein synthesis is still under debate. If we long thought that RPs were mostly required to ensure the structural integrity of the ribosome, several evidence tend to demonstrate that RPs can directly impact the extent of protein synthesis. Indeed, RPS6 phosphorylation has been proposed to regulate global protein synthesis, with a direct impact on translation initiation and elongation [25, 30, 31]. Originally, RPS6 phosphorylation has been thought to be important for specific translation, through the translational regulation of 5’-terminal oligopyrimidine mRNAs (5’-TOP mRNA) [32]. However this hypothesis has been rule out [23, 25]. A recent study comparing wild-type MEF and unphosphorylable RPS6 (RPS6^p-/p-^) MEF also found no difference in the level of 5’-TOP mRNA translation[33].

Interestingly, RPS6 phophorylation has been shown to be important for ribosome biogenesis [34]. In particular, RPS6 is involved in the transcriptional regulation of Ribosome Biogenesis (RiBi) factors involved in pre-rRNA synthesis, cleavage, post-transcriptional modifications, ribosome assembly and export. Therefore, one can hypothesize that increase of RPS6 phosphorylation promotes ribosome biogenesis and subsequent enrichment of the pool of ribosomes in cells. Axon regeneration requires extensive protein synthesis to generate all the building blocks to sustain axon growth, as illustrated by the level of global protein synthesis that is decreased in neurons upon axon injury [26]. Thus, increasing the number of ribosomes may help to sustain high levels of protein synthesis to support axon regeneration. However, RPS6 phosphorylation-mediated control of protein synthesis is still under debate, and may be cell type-dependent.

Furthermore, Bohlen and colleagues have demonstrated that the level of RPS6 phosphorylation differentially affects mRNA translation based on the length of their coding sequence (CDS) [33]. As ribosomes translate mRNAs, RPS6 are progressively dephosphorylated. Therefore, mRNA with long CDS harbor less phosphorylated RPS6. In contrast, mRNA with short CDS are actively translated by phosphorylated RPS6. Thus, RPS6 phosphorylation level may indicate which cell program is preferentially translated in neurons and could participate directly in the regenerative capacities observed in DRG. Finally, beyond the quantitative aspect of how many ribosomes cells have, the post-translational state of RP may be equally important in protein synthesis efficiency. In the context of axon regeneration, phosphorylation of RPS6 specifically at Ser235-236 appears to be critical to induce axon growth. Indeed, many genes involved in axon regeneration have short CDS and can be preferentially translated by ribosomes with phosphorylated RPS6 [33], such as ATF3 [35] (ATF3_MOUSE 181 aa), KLF family [36] (KLF7_MOUSE 301 aa), Rheb [17] (RHEB_MOUSE 184 aa), or genes implicated in mitochondria function [37] (PPARG_MOUSE 505aa, UCP2_MOUSE 309 aa, ARMX1_MOUSE 456 aa). Based on these observations, RPS6 phosphorylation may prime neurons for regeneration by facilitating the translation of pro-regenerative mRNAs.

We show that in DRG, RSK2 controls RPS6 phosphorylation, which in turn controls the preconditioning effect and axon regeneration both in the CNS and the PNS. RSK acts downstream of the MAPK pathway as it is activated by ERK1/2. The RSK family is closely related to the MSK (mitogen and stress activated kinase-MSK1 and 2) family. However, their physiological functions are different [38]. RSK have two kinase domains. The N-terminal kinase domain is an AGC family kinase that shares 57% of amino acids with the S6K1 kinase domain. The C-terminal kinase domain is related to the CAM-K kinase family. Thus, despite potential sharing of substrates with S6K1, RSK may have specific targets. RSK promotes the phosphorylation of RPS6 on Ser235-236, which in turn promotes the assembly of the translation complex. This process correlates with an increase of CAP dependent translation, independently of the mTOR pathway [39]. Our results show that RSK2/RPS6 phosphorylation-controls regeneration independently of mTOR activation. In fact, this suggests that mTOR and RSK2 will have different regenerative outcomes, possibly depending on the neuronal subpopulation. Importantly, in DRG, mTOR and RSK pathways are not redundant and they do not compensate each other.

In this study, we focused on the RSK-RSP6 axis, yet RSK is known to phosphorylate several other substrates that could participate in axon regeneration. For example, RSK2 controls the transcription regulation of c-fos [40] and CREB [41], which are both involved in axon regeneration [42]. RSK2 is also important in the process of cell growth based on its regulation of GSK3β phosphorylation [43]. Interestingly, GSK3 has been described to promote axon regeneration in CNS via its activity on MAP1D and CRMP2 [44]. Finally, RSK2 positively regulates cell survival. This activity may depend on regulation of transcription (for example through the control of CREB expression), but also on the direct phosphorylation of the protein Bad to further inhibits its pro-apoptotic effect [45]. As neuronal survival is key for the outcome of injury response in the PNS and the CNS, RSK2 may be implicated in the neuroprotection observed after sciatic nerve lesion. Altogether, a larger analysis of the diverse phosphorylated targets of RSK2 in DRG neurons and in CNS neurons will give us more insight into the precise mechanisms of RSK2-dependent regeneration.

Interestingly, at the time of preparation of the present work, a study from Mao and colleagues also identified RSK family as key effectors of PNS and CNS regeneration [46]. They found that RSK1 is upregulated in DRG neurons after sciatic nerve injury. Inhibition of RSK1 decreases sciatic nerve regeneration, whereas its overexpression enhances sciatic nerve regeneration. Mechanistically, authors described that overexpression of the elongation factor eEF2 rescues the effect of RSK1 inhibition both *in vitro* and *in vivo*. They showed that eEF2 controls the translation of pro-regenerative mRNAs such as BDNF or IGF, which have been largely described to modulate axon regeneration and neuroprotection [9, 47]. Both molecules partially rescue the deletion of RSK1 *in vitro*. Interestingly, based on their study and ours, RSK1 and RSK2 seem to have a similar pro-regenerative effects in the PNS. However, while both mechanisms of action are based on translational control, the modalities and effectors are different. eEF2 factor is a canonical translational factor implicated in the regulation of translation elongation. RSK1-mediated phosphorylation of eEF2 kinase, which controls eEF2 phosphorylation, leads to its inactivation and thus to the activation of eEF2 to promote global and specific translation necessary for regeneration. Both Mao and colleagues’ work and ours demonstrate that RSK family critically regulates the post-translational modification of components of the translational complex, thereby controlling protein synthesis and axon regeneration. To note, RSK2 can phosphorylate eEF2K and RSK1 can also phosphorylate RPS6. It would be interesting to decipher if RSK1 and 2 co-expression synergies to further enhance axon regeneration.

Mao and colleagues also tested the role of RSK1 in CNS regeneration. To analyze this feature, they use the visual system model, in which they overexpressed RSK1 in RGC by AAV2 intravitreal injection. RGC neurons cannot regenerate spontaneously after a lesion and, in contrast to DRG, injury signals induce neuronal death and suppresses growth capacities [10]. They showed that RSK1 expression level is not regulated by the optic nerve crush. Moreover, RSK1 overexpression alone has no effect on regeneration nor on neuroprotection in the visual system. This contrasts with our data showing that RSK2 can promote CNS regeneration of the dorsal column. Two hypotheses can explain this discrepancy. First, even if RSK1 and 2 share common targets and are often implicated in the same biological functions [24], RSK1- and RSK2-controlled targets are not strictly overlapping in cells, thus there is a specificity of action of each isoform. Second, there may be a cell type-specificity of action for the RSK family. Even if large spectrum of neuroprotective and regenerative molecular pathways are shared between the different neuronal populations of the CNS and PNS, neurons have cell type- and subpopulation-specific injury responses. For example, ATF3 upregulation in DRG following sciatic nerve injury promotes a positive response to injury, whereas injury-induced upregulation of ATF3 in RGC leads to neuronal death and inhibition of axon outgrowth. Besides, differential injury response of individual neuronal subpopulations has been recently addressed in RGC. sc-RNAseq analysis of single RGC allowed to highlight intrinsically-resilient subpopulations of RGC [48, 49]. This suggests that each subpopulation of neurons has an intrinsic specific machinery that influences its response to stress. In our case, the regenerative effect of RSK2 in other CNS regeneration models remains to be determined. Nonetheless, we can propose that DRG are more prompt to respond to RSK activity compared to RGC. Interestingly, PTEN deletion-induced activation of mTOR combined with RSK1 overexpression induces a synergistic effect on optic nerve regeneration, with more axons reaching the distal part of the optic nerve compared to PTEN deletion alone [46]. In DRG, we found that RSK2-mediated phosphorylation of RPS6 is mTOR-independent, whereas in RGC, mTOR may be required to phosphorylate RPS6, along with RSK1-mediated control of eEF2 activity. Altogether, mTOR-RSK interactions may well depend on the neuron type in order to control RPS6 phosphorylation.

To conclude, our work demonstrates that RPS6 phosphorylation is key in the process of PNS and CNS regeneration. Its regulation by RSK2 independently of mTOR highlights the role of this pathway in regeneration and open new avenues to understand molecular mechanisms of axon regrowth and functional recovery.

## Material and Methods

### Surgical procedures

Animal care and procedures were performed according to the Grenoble Institute Neurosciences, French and European guidelines. For intrathecal injections and dorsal column crush, mice were anesthetized with a mix of Ketamine (100mg/kg) and Xylazine (10mg/kg). Sciatic nerve crush procedure was performed under 3% induction, 2% maintenance isoflurane. For analgesia, Paracetamol was given in the drinking water (4mg/ml) one day before and 2 days after surgery. Buprenorphine (0.05mg/kg) was administered by subcutaneous injection 6h before dorsal column injury and every 6h for 3 days after surgery.

#### Animals

Mice with mixed backgrounds were used as wildtype animals, regardless of their sex. Phospho-dead RPS6 mouse line has been already described [25] and is maintained in mixed background. Animals were kept on a 12h light/dark cycle with food and water provided at libitum, at a constant temperature and humidity (21°C; 10% humidity). In all experiments, mice showing any signs of hindlimb paralysis or any discomfort were removed from further experiments.

#### AAV8 Virus Injections

3 to 4 weeks old mice were injected intrathecally, as described previously [50] using a 30G needle with 10uL of the following viruses: AAV8-PLAP (placental alkaline phosphatase; as control), AAV8-RSK2, AAV8-RSK3, AAV8-Sh-Scrambled or AAV8-Sh-RSK2. Virus titers were around 1×10^14 particles/mL

#### Sciatic nerve crush

The left sciatic nerve was exposed and crushed for 15 seconds with forceps (Fine Science Tools-Dumont SS Forceps) with an angle of 45°. The sciatic nerve was crushed again at the same place for 5 seconds to ensure that all axons have been axotomized.

#### Dorsal column injury

5-6 weeks old mice underwent laminectomy at the level of T7 vertebra exposing the spinal cord. Using modified fine forceps (Fine Science Tools; Dumont #5SF Forceps; maximum amplitude of 1mm, adjusted with a rubber), the dorsal column was crushed 3×10secs with 700um of depth.

#### Dorsal column retrograde labelling

After exposing the injured sciatic nerve, 3ul of CTB (Cholera Toxin Subunit B, Alexa Fluor™ 555 Conjugate-1mg/mL) was injected upper to the lesion site with a glass micropipette to analyze the extend of dorsal column regeneration in the spinal cord.

### Adult DRG neurons culture, drug treatment and immunostaining

DRG cultures were performed as describes previously [51]. DRG from WT or animals that underwent intrathecal injection of AAV8 two weeks before were collected. Briefly, lumbar DRG (L3-L4 and L5) were dissected out and collected in iced cold Hank’s balanced salt solution (HBSS; Gibco). DRG tissues were incubated in 5% Collagenase A (Roche) at 37°C for 70mins and in 0.25% Trypsin (Gibco) for 5min. DRG were gently dissociated with blunt glass pipettes. Neurons were purified on a 10% bovin serum albumin (sigma Aldrich) cushion and plated on Poly-L-lysine (10mg/mL Sigma Aldrich) and Laminin (0,5mg/mL-Sigma Aldrich) coated cover slips in Neurobasal A medium (Gibco) supplemented with 2% B-27 supplement (Gibco) and 1% L-glutamine (Gibco). Cultures were maintained in a humidified atmosphere at 5% CO2 in air at 37°C.

In case of drug treatment, neurons were treated 1h after plating with RSK inhibitor BRD7389 at 3uM (Santa Cruz), mTOR inhibitors Torin1 5nM (Santa Cruz) or Rapamycin 0.1nM (Sigma Aldrich), S6K1 inhibitor PF-4708671 8uM (Sigma Aldrich) and Translation inhibitor Cycloheximide 2nM (Sigma Aldrich). All the dilutions were performed in DMSO. For each group treated with drugs, the respective control received DMSO treatment.

After 16h of culture, neurons were fixed in PFA 4%/sucrose 1.5% in PBS, permeabilized in 0,5% Triton (Sigma Aldrich) and labeled with primary antibodies, anti-Tuj1 (1/500, Biolegend), anti-pRPS6 (Ser 235-236; 1/500-Cell signaling Technology), anti-pRPS6 (Ser 240-244; 1/500-Cell signaling Technology), anti-RSK (1/500, RSK123-Cell signaling Technology) for 2h at room temperature and secondary antibodies, Alexa-488, Alexa-647, and Alexa-568 (1/800-ThermoFisher) for 1h at room temperature. Coverslips were mounted with Fluoromount-G™ Mounting Medium, with DAPI (Invitrogen).

### Cell transfection and protein extraction

N2A cells were transfected using PEI (3mg/mL; Alfa Aesar-ThermoFisher) with 3ug of the following plasmids: pAAV-vsvg-RSK1, pAAV-flag-RSK2, pAAV-v5-RSK3, pAAV-his-RSK4, pAAV-PLAP, pAAV-CMV-mCherry-U6-sgRNA_shRSK2 (5’ GGACCAACTACCACAATACCA 3’), pAAV-CMV-mCherry-U6-sgRNA_shCtrl (5’ GCAACAACGCCACCATAAACT 3’). These plasmids were obtained by cloning cDNA extracted from mouse cerebellum in pAAV-MCS Expression Vector with In-Fusion Cloning system (Takara). 96h after transfection, proteins were extracted using RIPA (Triton 1%) buffer supplemented with protease (Roche) and phosphatase inhibitors (Roche).

For DRG, proteins were extracted using 10mM Tris-HCl pH 7.5; 150mM NaCl; 0.5mM EDTA; 1% NP-40 with protease and phosphatase inhibitors (Roche). DRG were further lysate by sonicated (Vibra-Cell™, VWR) 5 times 10 secs.

Proteins were quantified with BCA following manufacturer’s instructions (Pierce- ThermoFisher).

### Westernblot

20ug of proteins from N2A cells or 10ug of protein from DRG were loaded to SDS-polyacrylamide precast gels (Biorad) and transferred to nitrocellulose membranes. Membranes were stained with Ponceau Red to verify the quality of the gels. Membranes were the blocked in 5% low fat milk for 1h in Tris Buffered Saline with 0.05% of Tween-20 (TBST) for 1h at room temperature and incubated over night at 4°C with the following antibodies diluted in blocking solution: anti-RSK2 (1/1000, Cell Signaling Technology); anti-RPS6 (1/1000, Cell Signaling Technology), anti-phospho-RPS6 ser235/236 (1/1000, Cell Signaling Technology), anti-phospho-RPS6 ser240/244 (1/1000, Cell Signaling Technology), anti-Flag (1/2000, Sigma Aldrich), Anti-His (1/5000, Proteintech), anti-RFP (1/1000, Abcam); anti-GFP (1/1000, Abcam) or GAPDH (1/5000, Proteintech). Membranes were then incubated for 2h at room temperature with HRP-coupled secondary antibodies (anti-rabbit HRP, Proteintech; anti-mouse HRP, Thermofisher Scientific) diluted from 1/1000 to 1/10000 in blocking solution. Membrane were developed with ECL (1.5mM luminol, 0.225mM coumaric acid, 100mM Tris HCl, 0.1mM hydrogen peroxide in miliQ water) using a chemidoc (ChemiDoc MP, Biorad).

### Histological procedures

After intracardiac perfusion of mice with iced cold PFA-4%, tissues were dissected-out and post-fixed in 4% PFA for 2h for sciatic nerve and DRG, overnight for spinal cords at 4°C. After cryopreservation in 30% sucrose, tissues were sectioned using a cryostat: DRG and sciatic nerves were sectioned at 12um and spinal cords at 20um.

#### Immunohistochemistry

Sections were blocked in blocking solution (5% BSA, 1% Donkey Serum, 0.5% Triton in DPBS) for at least 1h at room temperature. Sections were then incubated overnight at 4°C with primary antibodies diluted in blocking solution: anti-βIII Tubulin (1/500, Biolegend), anti-RSK2 (1/100, Cell Signaling Technology), anti-RPS6 (1/100, Cell Signaling Technology), anti-phospho-RPS6 ser235/236 (1/100, Cell Signaling Technology), anti-phospho-RPS6 ser240/244 (1/500, Cell Signaling Technology), Anti-His (1/1000, Proteintech), anti-RFP (1/500, Abcam); anti-Flag (1/1000, Sigma), anti-GFP (1/400, Abcam), anti-GFAP (1/500, ThermoFisher), anti-SCG10 (1/1000, Novus) or anti-CTB (1/500, Abcam). Then tissues were incubated with the appropriated secondary antibodies (Alexa-fluor conjugated-Jackson laboratories) diluted in blocking solution at 1/500 for 2h at room temperature. Slides were mounted with Fluoromount-G™ Mounting Medium, with DAPI Medium (Invitrogen).

#### *In situ* hybridization

Experiments were performed as described in Nawabi et al 2010 [52]. Probes were cloned in pGEMT vector from cDNA extracted from cerebellum: pGEM-T_RNAprobeRSK1; pGEM-T_RNAprobeRSK2; pGEM-T_RNAprobeRSK3; pGEM-T_RNAprobeRSK4 Sequence used for the probe was describe in Tab.S1.

### Image Analysis and quantification

DRG cultures, DRG sections, sciatic nerves sections were imaged using a Nikon Ti2 Eclipse epifluorescent microscope with 4x; 10x and 20x objectives. Spinal cord sections were imaged using a Dragonfly spinning disk microscope (Andor Technologies) with a 20x objective. Sections were stitched using the Fusion software with 10% overlap between tiles.

#### Analysis of neurite outgrowth, ramification and survival

The mean of neurite outgrowth, ramification and survival of DRG neurons was manually measured with ImageJ software. The mean neurite outgrowth for at least 50 neurons per condition (except for BRD7389 and cycloheximide condition) was quantified for at least at 3 independent biological replicates. Neurite ramification was analyzed for at least 30 neurons per condition from at least at 3 independent biological replicates. DRG neurons survival was quantified from 10 random microscope fields per condition from at least 3 independent biological replicates.

#### Analysis of sciatic nerve regeneration

Axon regeneration was quantified on 3 to 5 sagittal sections for each mouse. SCG10 intensity was quantified with ImageJ software in 500um area, the background intensity was subtracted and residual intensity was compared to the crushed area (maximum intensity of SCG10).

#### Analysis of dorsal column regeneration

Axon regeneration was quantified on 2-4 sections for each mouse. CTB intensity was quantified with ImageJ software every 50um, the background intensity was subtracted and residual intensity was compared to maximum intensity. The origin of regeneration/dieback (distance “0”) is the caudal part of the crush site.

### Behavior tests

For behavior tests, we used mix background, male and female mice from pooled litters. Before the first surgery (intrathecal injection), mice were handled once per day with soft and strong contention, head belly and foot contact. After the first surgery, for the Von frey filament, mice were placed 10min per day during 7 days in a 10cm diameter glass ramekin on non-sharpness grid at 60cm above the floor. For the removal of the sticky paper, mice were placed on individual cages and trained 7days on active phase with the sticky paper stuck in both paws until they were able to remove the sticker. After training, all experiments were performed once a week, 2 weeks before dorsal column injury and 6 after.

#### Sticky paper attention

For this test, mice were placed in the experiment room at least 1h before behavior test and the experiment is performed during the activity period of mice (during night) with red light only. Mice were placed in a transparent “look-like” home cage. After at least 5min of acclimatization, an 8mm diameter adhesive pad was stuck to each hind paw. Time of first contact between mice nose and the sticky paper and the time needed for its removal was quantified for each hind paw. This experiment was performed twice each time with a maximum given time of 5min to remove both pad [53]. Mice activity was recorded with two cameras (Logitech HD 1080p/ 60 fps) for further analyses.

#### Von Frey filament test

For this test, animal was place in the experiment room at least 1h before behavior. Each mouse was individually placed in a 10cm diameter bottomless box 10min before the test. The box was placed on non-sharpness grid at 60cm above the floor. The method used in this test is the up and down method as described previously [54]. Briefly, once mice had calm down, they were tested for 3 seconds with the reference filament in the center of the paw. In case of reaction, the next test was performed with smaller filament (more sensitive). If the mice do not response, the next test was performed with a thicker filament (less sensitive). To determine mice sensitivity, they should respond three times for the same filament. This experiment was done for both paws independently. Mice activity was recorder with camera (Logitech HD 1080p/ 60 fps) for further analyses.

### Statistical analysis

All animals used are both male and females from pooled litters and were randomly assigned to groups before any treatment or experimental manipulation. All analyses were performed while blinded to the control test realized in same time. Statistical analysis was performed with GraphPad Prism 9.4 using either One sample t test, One-way ANOVA, Two-way ANOVA, Kruskal wallis test; Paired t-test, Unpaired t-test. Each test used is indicated in figure legends. Error bars indicate the standard error of the mean (SEM). A *p*-value p < 0.05 was considered statistically significant with difference indicate by stars: ⁎⁎⁎ p<0.001, ⁎⁎ p<0.01, ⁎ p<0.05.

## Author Contribution

**Conceptualization:** Charlotte Decourt, Homaira Nawabi, Stephane Belin

**Data curation**: Charlotte Decourt, Julia Schaeffer, Homaira Nawabi, Stephane Belin

**Formal analysis**: Charlotte Decourt, Stephane Belin

**Funding Acquisition**: Homaira Nawabi, Stephane Belin

**Investigation**: Charlotte Decourt, Julia Schaeffer, Beatrice Blot, Blandine Excoffier, Antoine Paccard, Homaira Nawabi, Stephane Belin

**Methodology**: Charlotte Decourt, Mario Pende, Stephane Belin

**Project administration**: Homaira Nawabi, Stephane Belin

**Ressources**: Mario Pende, Stephane Belin

**Supervision**: Charlotte Decourt, Homaira Nawabi, Stephane Belin

**Visualization**: Charlotte Decourt, Homaira Nawabi, Stephane Belin

**Writing original draft**: Charlotte Decourt, Homaira Nawabi, Stephane Belin

**Writing review and editing**: Charlotte Decourt, Julia Schaeffer, Mario Pende, Homaira Nawabi, Stephane Belin

## Acknowledgments

We would like to acknowledge Pr. H. Zhigang for his input and E.Plissonier; T.-N.Nguyen; N.Fayad, A.Lapierre for laboratory help and discussion. We thank S. Carnicella, M. Bartolomucci and the GIN behavioral facility that is supported by the Grenoble Center of Excellence in Neurodegeneration (GREEN). This work was supported by the Photonic Imaging Center of Grenoble Institute Neuroscience (Univ Grenoble Alpes – Inserm U1216) which is part of the ISdV core *facility and* certified by the *IBiSA* label

---

**S1 Fig.**
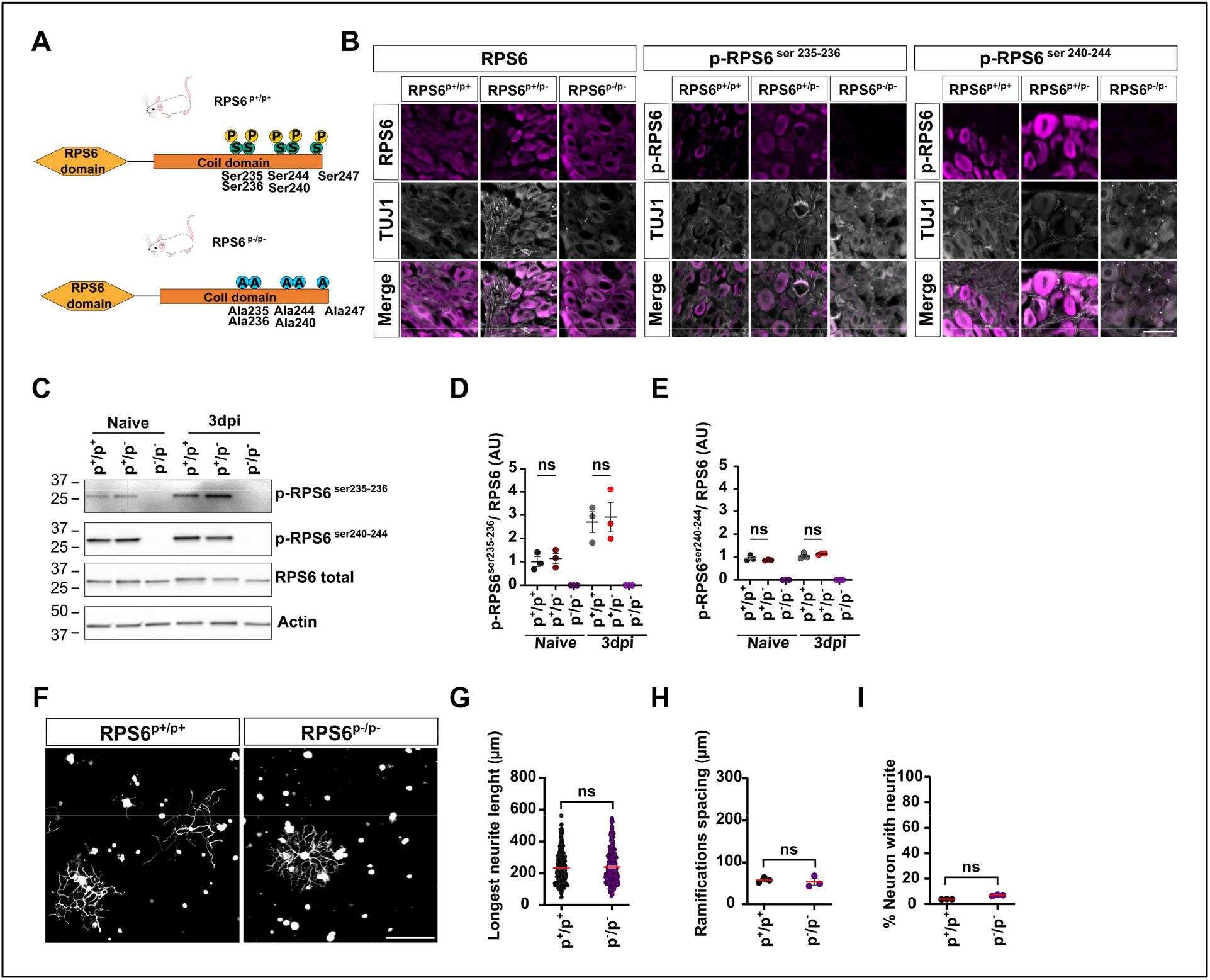
Characterization of phosphor-dead RSP6 mouse line. (A) Schematic describing the phospho-dead RSP6 mouse line. (B) Western blot showing that RSP6 is not phosphorylated in RPS6^p-/p-^ DRG compared to RPS6^p+/p+^ RPS6^p-/p+^ DRG. (C-D) Quantification of B. (Mean ± SEM; One sample t test; N=3 independent animal/groups) (E) Representative microphotographs of DRG sections stained with anti-RPS6; anti-p-RSP6^Ser235-236^ or anti-Phospho-RPS6^ser240-244^ (in magenta) and anti-Tuj 1 (in gray) scale bar: 25um. (F) Representative microphotographs of naive cultures of mature DRG neurons from WT (RPS6^p+/p+^) and homozygous (RPS6^p-/p-^) mice line defective for RPS6 phosphorylation showing no differences. Scale bar: 250um (G-I) Graphs showing the quantification of D. (G) Longest neurite after per neuron 16h after plating (Mean ± SEM; Unpaired t test with Welch’s correction One-way ANOVA, Dunn’s multiple comparisons test, 3 independent DRG cultures, approximately 50 cells per conditions per culture). (H) Distance between two ramifications in longest neurite were analyzed; Mean ± SEM Unpaired t test; 3 independent DRG cultures; approximately 50 cells were analzed per conditions per culture. (I) Percentage of neurons with a neurite 16h after plating Mean ± SEM Unpaired t test; 10 random microscopy fields were quantified per condition). ns: non-significant).

**S2 Fig.**
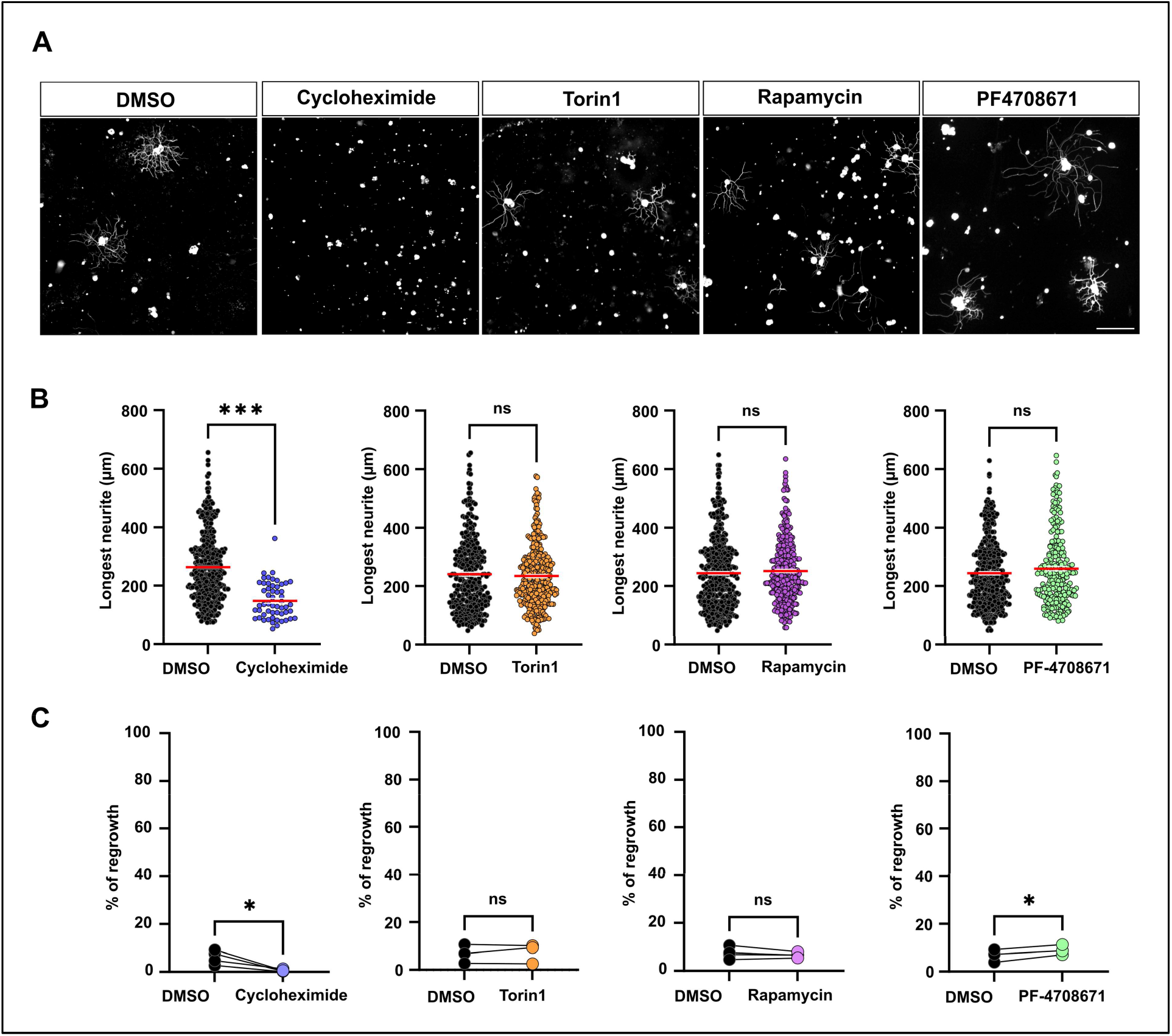
Characterization of the effect of different signaling pathways on naive DRG cultures. (A) Representative microphotographs of naive DRG neurons cultures treated with DMSO (control), a global protein translation inhibitor (cycloheximide (5nM)); mTOR inhibitors (Torin (5nM) or Rapamycin (0.1nM)), and a S6K1 inhibitor (PF-4708671 (8uM)). Scale bar: 250 um. (B) Quantification of A (Mean ± SEM; Kruskal wallis test Dunn’s multiple comparisons test; 3-4 independent DRG cultures; approximately 50-100 cells were counted per condition per culture (except for cyclohexamide) (C) Percentage of neurons with a neurite 16h after plating from A (Mean, Paired t-test. 3-4 independent DRG cultures; 10 random microscopy fields were quantified per condition) ⁎⁎⁎ p<0.001, ⁎⁎ p<0.01, ⁎ p<0.05.

**S3 Fig.**
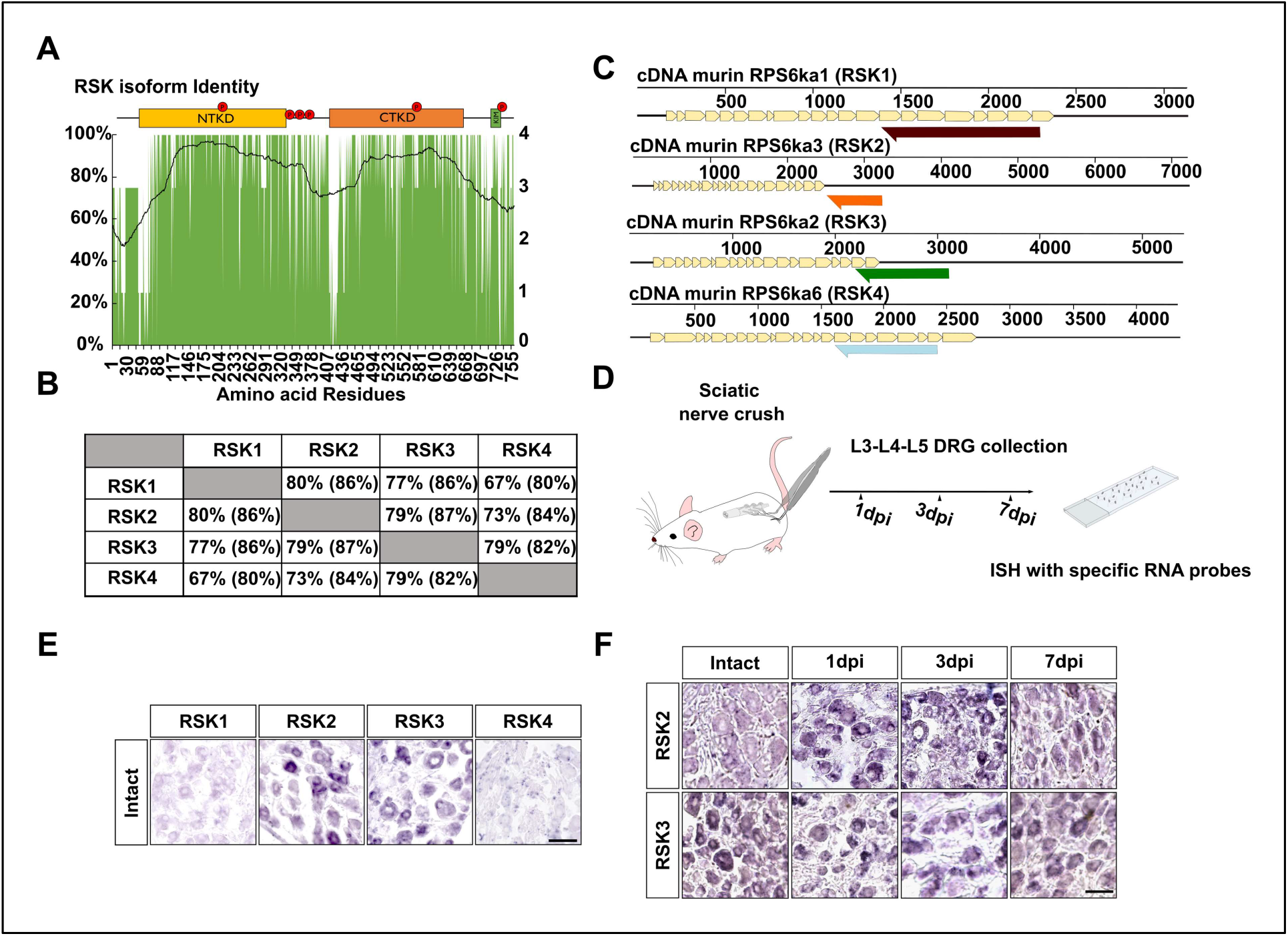
Characterization of RSK family expression in mature DRG. (A) Graph showing the homology of amino-acid sequence among the 4 RSK expressed in mouse. (B) Table summarizing the homology and identity among RSK1, 2, 3 and 4. (C) Schematic of the probes used to study specific expression of RSK1, 2, 3 and 4 by *in situ* hybridization. (D) Workflow of experiment. (E) Microphotographs showing *in situ* hybridization of RSK1, RSK2, RSK3 and RSK4 on adult lumbar DRG sections. Only RSK2 and RSK3 are highly expressed in mouse lumbar DRG. (F) Microphotographs of DRG sections showing *in situ* hybridization of RSK2 and RSK3 on intact and after sciatic injury at 1, 3 and 7-days post injury (dpi). RSK2 expression is regulated by axon injury. Scale bar: 50um.

**S4 Fig.**
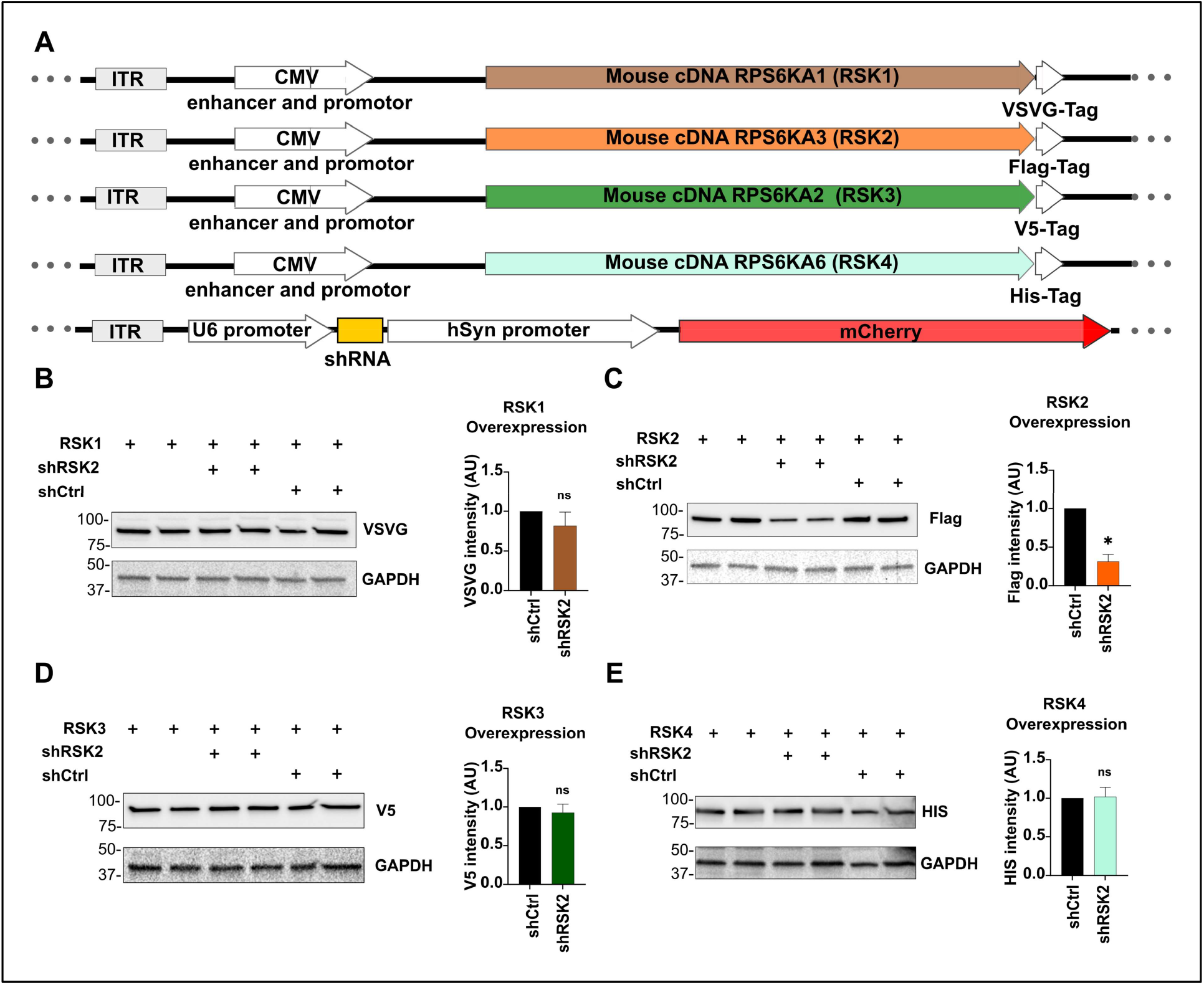
Characterization of ShRNA-RSK2. (A) Schematic of the plasmid constructs used to overexpress RSK1-vsvg, RSK2-flag, RSK3-v5, RSK4-his, PLAP or ShRNA (Sh-scrambled or Sh-RSK2). (B-E) Western blot showing ShRNA-RSK2 specificity in N2A cells 96h after co-transfection. (Mean ± SEM; One sample t-test; N=3 individual transfections per group. ⁎⁎⁎ p<0.001, ⁎⁎ p<0.01, ⁎ p<0.05.

**S5 Fig.**
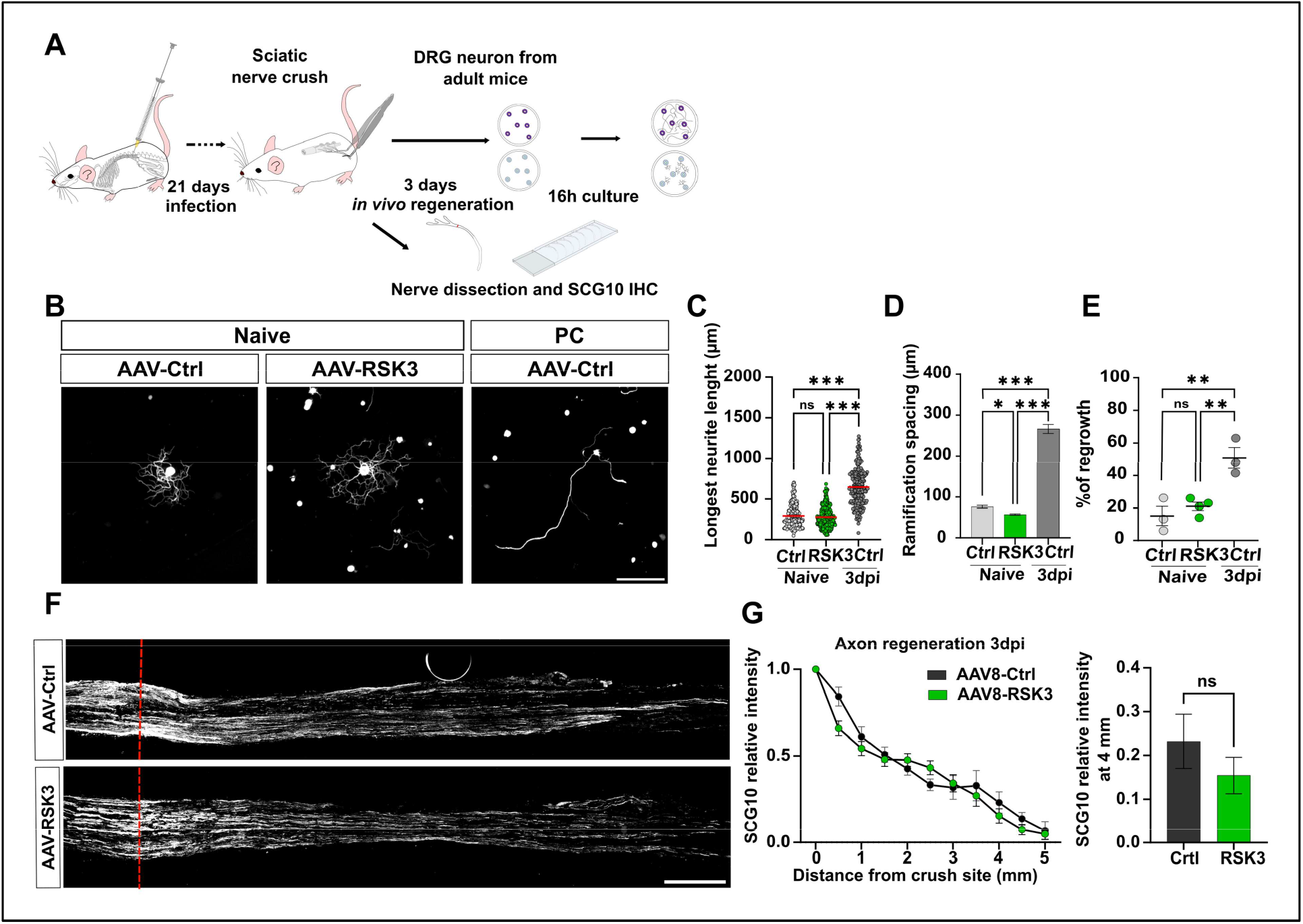
RSK3 is not involved in the preconditioning effect. (A) Representative microphotographs of DRG dissociated cultures showing that RSK3 overexpression in naive cultures does not phenocopy the preconditioning effect. Scale bar: 250um (B-D) Quantification of A. (B) Longest neurite per neuron 16h after plating (Mean ± SEM One-way ANOVA Dunn’s multiple comparisons test, at least 3 independent DRG cultures, approximately 50-100 cells were counted per condition per culture); (C) Mean distance between two ramifications (Mean ± SEM Two-way ANOVA Tukey’s multiple comparisons test; at least 3 independent DRG cultures; approximately 50 cells per condition per culture) and (D) Percentage of neurons with a neurite 16h after plating (Mean ± SEM Two-way ANOVA Tukey’s multiple comparisons test; 10 random microscopy fields quantified per conditions per culture). (C) Representative confocal images of the sciatic nerve sections 3 days post-injury from mice injected intrathecally with AAV8-PLAP (control) or AAV8-RSK3. Regenerating axons are labeled with anti-SCG10 antibody (white). Red dashed line indicates the injury site. Scale bar: 500 um. (D) Quantification of regenerative axons from D (Mean ± SEM Multiple Unpaired t-test At least 3 independent animals per group) ⁎⁎⁎ p<0.001, ⁎⁎ p<0.01, ⁎ p<0.05.

**S6 Fig.**
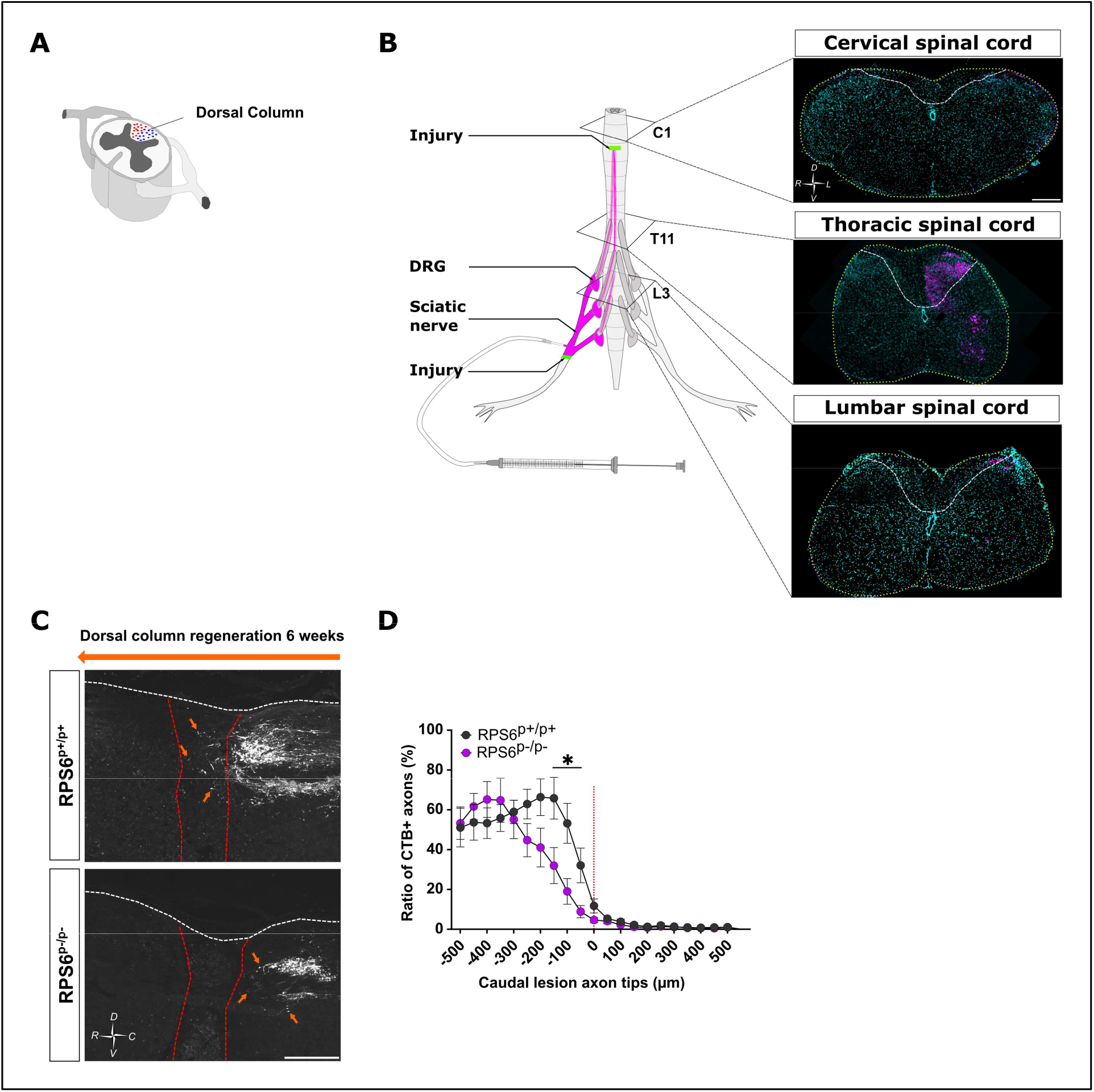
RPS6 phosphorylation is necessary for dorsal column regeneration in preconditioned condition. (A) Schematic representation of the dorsal column. (B) Representative images of cervical, thoracic and lumbar coronal sections of mice 6 weeks after dorsal column crush at thoracic T7 level, 1 week after CTB-Alex-555 intranervous injection in the sciatic nerve. (C) Representative confocal images of thoracic spinal cord sagittal sections 6 weeks after sciatic nerve and dorsal column crush from RPS6^p+/p+^ or RPS6^p-/p-^ mice. Regenerative axons are labeled with anti-CTB antibody (white). (D) Quantification of axon regeneration and dieback from caudal marge of crush site from B. (Mean ± SEM; Mann-Whitney test; N= at least 6 independent animals per condition).

**S1 Table. List of cDNAs used for *in situ* hybridization.**

